# Evaluating spatially variable gene detection methods for spatial transcriptomics data

**DOI:** 10.1101/2022.11.23.517747

**Authors:** Carissa Chen, Hani Jieun Kim, Pengyi Yang

## Abstract

The identification of genes that vary across spatial domains in tissues and cells is an essential step for spatial transcriptomics data analysis. Given the critical role it serves for downstream data interpretations, various methods for detecting spatially variable genes (SVGs) have been proposed. The availability of multiple methods for detecting SVGs bears questions such as whether different methods select a similar set of SVGs, how reliable is the reported statistical significance from each method, how accurate and robust is each method in terms of SVG detection, and how well the selected SVGs perform in downstream applications such as clustering of spatial domains. Besides these, practical considerations such as computational time and memory usage are also crucial for deciding which method to use. In this study, we address the above questions by systematically evaluating a panel of popular SVG detection methods on a large collection of spatial transcriptomics datasets, covering various tissue types, biotechnologies, and spatial resolutions. Our results shed light on the performance of each method from multiple aspects and highlight the discrepancy among different methods especially on calling statistically significant SVGs across datasets. Taken together, our work provides useful considerations for choosing methods for identifying SVGs and serves as a key reference for the future development of such methods.

## Introduction

Advances in spatial transcriptomics have made it possible to identify genes that vary across spatial domains in tissues and cells^1^. The detection of spatially variable genes (SVGs) is essential for capturing genes that carry biological signals and reducing the high-dimensionality of the spatial transcriptomics data^1^, which is akin to defining highly variable genes (HVGs)^2^ in single-cell RNA-sequencing (scRNA-seq) data^3^. These SVGs are therefore useful for various downstream analyses of spatial transcriptomics data. Spatially variable genes are conceptually different from HVGs found in scRNA-seq data as, by definition, SVGs preserve the spatial relationships of tissues and cells in the biological samples whereas HVGs do not necessarily preserve such relationships.

A fast-growing number of methods for SVG detection have been proposed in the recent literature. Some popular examples include SpatialDE^1^ based on Gaussian process; SPARK^4^ and SPARK-X^5^ based on mixed and non-parametric models, respectively; SOMDE based on self-organising map^6^; Giotto based on statistical enrichment of spatial network in neighbouring cells^7^; nnSVG based on nearest-neighbour Gaussian processes^8^; MERINGUE based on nearest-neighbour spatial auto-correlation ^9^, and Moran’s I as implemented in the Seurat package^10^. While various SVG detection methods have been incorporated into the typical workflows and pipelines for spatial transcriptomics data analysis such as the Giotto and Seurat packages, there is a lack of systematic evaluation and comparison of different methods. In particular, essential questions including the degree of agreement among different methods in terms of the ranking and selection of SVGs, the reproducibility of these methods in terms of SVG detection when the genes included in a given dataset changes, and the accuracy and robustness of SVG detection, and the utility of the selected SVGs to perform in downstream data analysis such as spatial domain clustering remain to be addressed. In addition, practical considerations such as running time and memory usage required by each method have not been systematically benchmarked.

To fill this critical gap, we systematically evaluated a panel of eight popular SVG detection methods on a collection of 31 spatial transcriptomics and synthetic spatial datasets. These datasets together capture various sample and tissue types and major spatial biotechnologies with different profiling resolutions, including Visium (10X Genomics), ST ^11^, Slide-seq ^12^, Slide-seqV2 ^13^, MERFISH ^14^, seqFISH+ ^15^, Stereo-seq ^16^, SM-Omics ^17^, and DBit-seq ^18^. Our results shed light on the performance of each tested SVG detection method in various aspects and highlight some of the discrepancies among different methods especially on calling statistically significant SVGs across datasets. Taken together, this work provides useful information for considering and choosing methods for identifying SVGs whilst also serving as a key reference for future development of SVG detection methods from spatial transcriptomics data.

## Results

### Evaluation framework and data summary

We designed an evaluation framework to gain insight into the performance of different SVG detection methods to call SVGs from a collection of real and simulated spatially resolved transcriptomics datasets (**Figure 1**). These include spatial transcriptomics data with varying sequencing depths generated from a wide range of spatial profiling platforms, species, tissue types, and spatial resolutions (**Supplementary Figure 1**). Specifically, our evaluation framework entailed a wide range of comparative and benchmarking analyses to investigate key questions. First, we compared the concordance between the overall rankings of the SVGs between SVG tools and evaluated their dependence on mean gene expression to assess the variability among methods and their capacity to account for the bias between gene expression and variance. Next, we investigated the capacity of each SVG method to reproducibly rank SVGs independently of the pool of genes observed in the dataset or with induced sparsity in spots, to call ground truth SVGs from synthetic spatial data, and to define SVGs required to accurately cluster spatial domains. Finally, using the spatial benchmarking datasets we compared the computational cost in terms of speed and memory required for SVGs to be called by each method.

**Figure 1.**
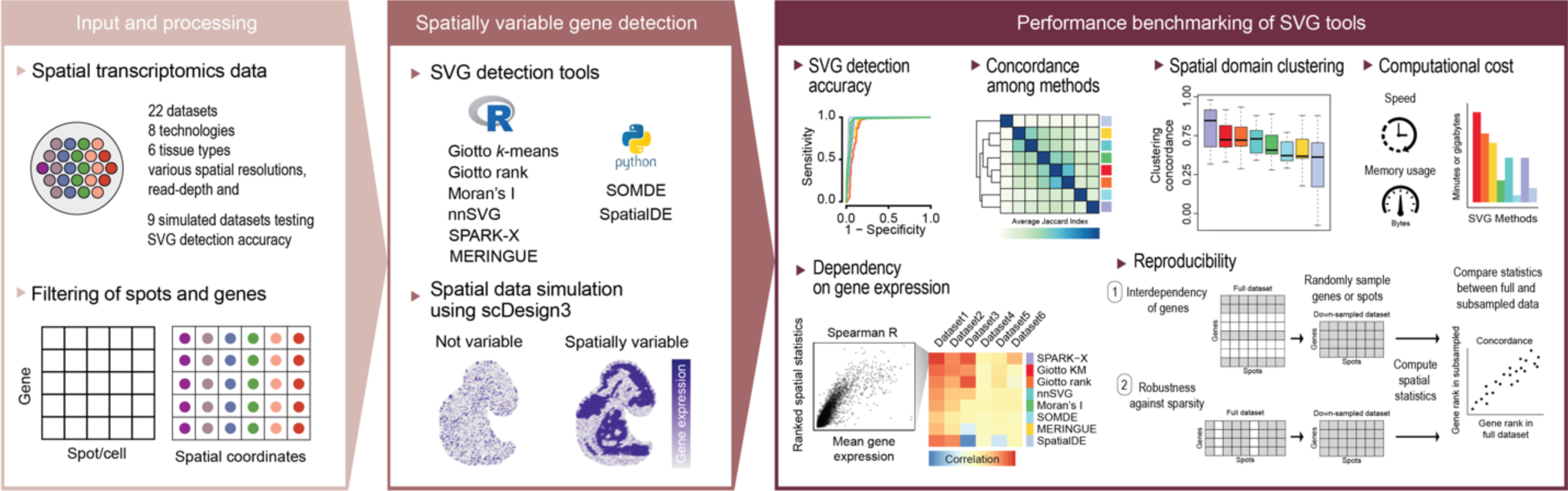
Schematic summary of the evaluation framework used in this study.

### Concordance among SVG detection methods

To quantify the degree of agreement amongst the different SVG detection methods, we first obtained the ranking of genes in each dataset ordered from the most to least spatially variable based on the statistics reported by each method and correlated the SVG rankings from each pair of methods. These correlation results were summarised for each SVG detection method with respect to other methods across the spatial datasets in **Figure 2a** and visualised individually in **Supplementary Figure 2.** The overall concordance results showed two groups of methods that highlighted an average similarity (measured as the Spearman’s correlation of SVG statistics) of greater than 0.8 across the spatial datasets. The most correlated pair of methods were Giotto K-means and Giotto rank, as expected, because of a large overlap in their framework to perform spatial network enrichment. The next group of correlated methods were MERINGUE, Moran’s I, and nnSVG. SOMDE, SPARK-X, and SpatialDE showed the least concordance with the other methods, suggesting the prioritisation of SVG statistics by these methods, in particular SpatialDE, are different to other methods. Amongst the methods, we observed that SpatialDE demonstrated the highest variability across datasets. Colouring the data points in Figure 2a by the total number of spatial spots and technology (**Supplementary Figure 3a-b**) revealed an interesting trend, which was most striking in SpatialDE, where despite an overall low correlation in spatial statistics with all other methods a high correlation was observed in specific datasets derived from the 10X Visium platform. Overall, these results demonstrate that whilst we observed moderate-to-high correlation between SVG detection tools in terms of SVG ranking, we found considerable variability of reported SVG statistics across the computational methods, platforms, and datasets.

**Figure 2.**
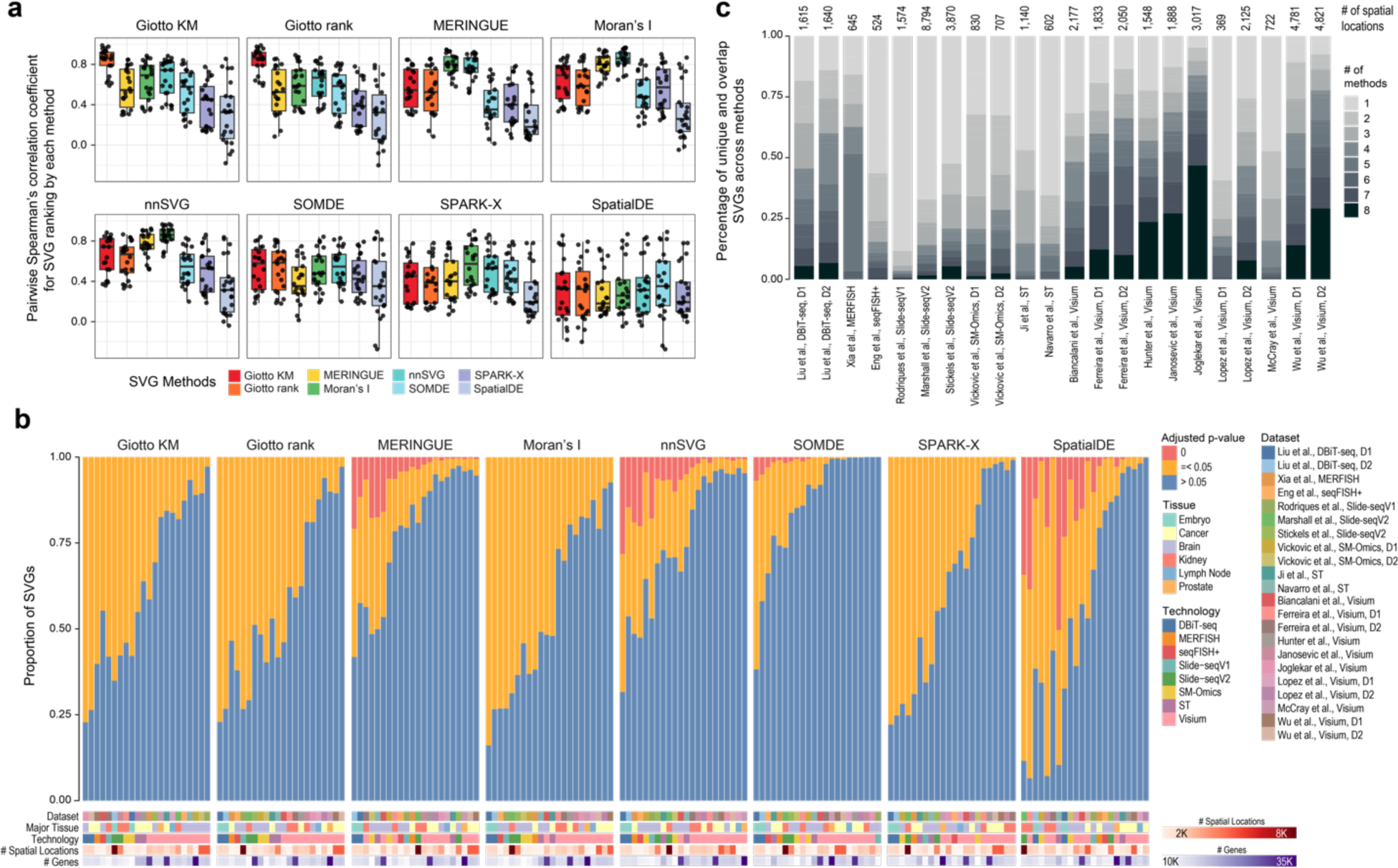
Concordance, statistical significance, and overlap of SVGs detected by different methods. (**a**) Concordance of SVG rankings reported from each SVG detection method. Each panel uses one SVG detection as an anchor and the y-axes are pairwise Spearman’s correlation coefficient for quantifying concordance in ranking of each pair of SVG detection methods. Points in each boxplot represent the result from a dataset. Boxplot centre line, median; box limits, upper and lower quartiles; whiskers, 1.5 times the interquartile range. (**b**) Statistical significance of SVGs detected by each method. SVGs are partitioned into three categories based on the adjusted p-values reported by each method (i.e., p = 0; 0 < p <= 0.05; p > 0.05) and presented as a percentage (y-axis). The datasets are ordered in terms of the decreasing proportion of genes observed in the orange category. The colour bars denote various characteristics of the spatial dataset including the tissue type, spatial technology, number of spatial locations, and total number of genes expressed. (**c**) A proportional bar plot showing the percentage of unique (# of method = 1) and overlapping (# of method > 1) significant SVGs (adjusted p-value <=0.05) reported by the SVG detection methods for each spatial transcriptomics dataset.

While the ranking of SVGs is useful for selecting the top candidates for subsequent analysis, in practice, statistical significance such as *p*-values is frequently used for selecting SVGs. To this end, we first partitioned the SVGs into three categories (i.e., *p* = 0; 0 < *p* <= 0.05; *p* > 0.05) based on the adjusted *p*-value reported from each computational method (**Figure 2b**). We found that most methods report a large proportion of SVGs at an adjusted *p*-value threshold of 0.05 on many datasets. Among the eight methods, nnSVG, MERINGUE, and SpatialDE, and to a lesser degree SOMDE, reported a sizable proportion of SVGs with an adjusted *p*-value of 0. Interestingly, SOMDE reported on average the fewest number of significant SVGs with some datasets having almost no significant SVGs (**Figure 2b and Supplementary Figure 3b**). Intriguingly, we observed that despite the high correlation in SVG statistics (**Figure 2a**), different methods predicted a vastly differing number of SVGs as significant using a p-value threshold of 0.05. However, we note that the overall pattern between methods is still similar when we compute the average concordance in gene sets of the top 200, 500, 1000, and all significant SVGs across all the datasets between methods (**Supplementary Figure 4**). As before, SpatialDE demonstrated the least similarity against all other methods, followed by SPARK-X and SOMDE (**Supplementary Figure 4**). Giotto KM and Giotto ranks again demonstrated a high similarity, but this time Moran’s I’s gene sets tended to show a higher concordance with the Giotto methods rather than MERINGUE and nnSVG, suggesting that whilst the overall ranking in gene statistics may be similar between Moran’s I and MERINGUE and nnSVG, the top most significant SVGs identified by Moran’s I appear to be more similar to those of the Giotto methods (**Supplementary Figure 4**). Importantly, despite the relatively high correlation in SVG statistics observed between methods, the number of SVGs found by all methods are strikingly low with many datasets having close to no overlapping SVGs across all eight computational methods (**Figure 2c**). In addition, many unique genes were found by various individual methods in most datasets (**Figure 2c**). Together, these findings highlight the discrepancy among methods when an adjusted *p*-value threshold of 0.05 was used for calling statistically significant SVGs.

### Dependency of SVG statistics on gene expression levels

In scRNA-seq data, it is known that variance in gene expression is positively correlated with gene expression level; therefore, most highly variable gene (HVG) detection methods implement procedures to account for this bias ^2^. To test whether methods designed for SVG detection have the tendency to select genes with higher expression levels, we investigated the correlation between mean gene expression and the SVG statistics for each method and dataset pair. We found that indeed the rankings of SVGs from most methods correlated positively with the mean gene expression (**Figure 3a, b**). In particular, SPARK-X showed average correlations of around 0.8 across the datasets (**Figure 3c**), and the Giotto methods and nnSVG showed correlations of around 0.5 across the datasets, suggesting a high dependency of SVG ranking on gene expression for these methods. We also correlated the proportion of zeros in gene expression across cells against SVG ranking for each method (**Supplementary Figure 5a, b**). Since the proportion of zeros is known to be negatively correlated with their expression levels, the negative correlation observed among each method and dataset pair further confirms the dependency we found between SVG ranking and gene expression among current SVG detection methods.

**Figure 3.**
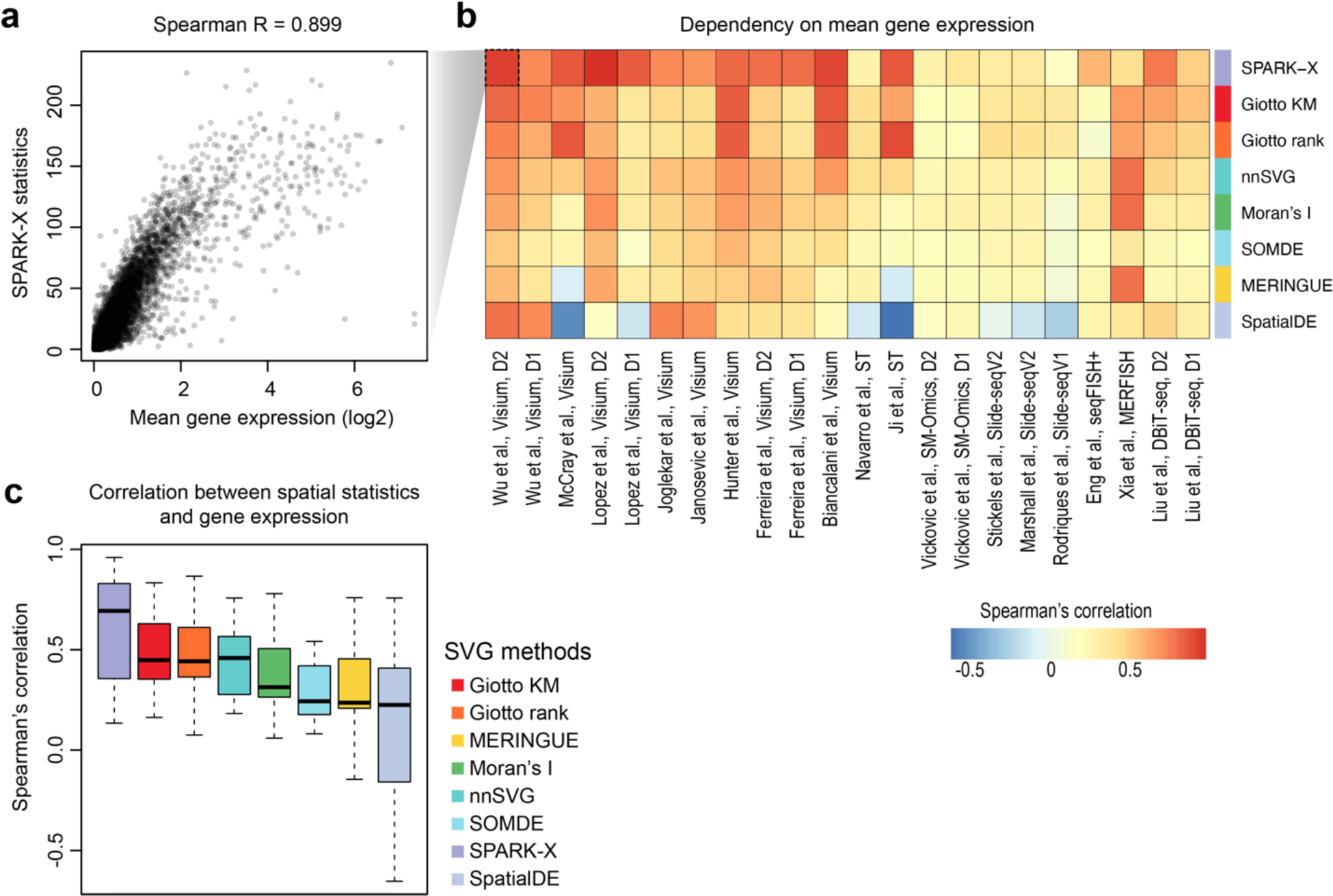
Dependency of SVG statistics on gene expression level. (**a**) An example showing the positive correlation between SVG statistics reported from SPARK-X and mean gene expression across cells in the “Wu et al., Visium D2” dataset^14^. (**b**) Heatmap summarising the Spearman’s correlation of SVG statistics reported by each method and the mean gene expression in each dataset. The rows are ordered from the highest to lowest average dependency. (**c**) Boxplot of Spearman’s correlation of SVG statistics reported by each method and the mean gene expression across the spatial transcriptomics datasets.

### Dependency of SVG statistics across genes and spatial spots

We next assessed the reproducibility of gene ranks based on the SVG statistics reported from each method when either the number of genes or the total number of spatial spots included in a dataset changes. To this end, we randomly down-sampled the genes in all benchmarking datasets (**Figure 4a**) to 50% and re-calculated the ranks of genes from the reported SVG statistics of each method on the down-sampled datasets. Most methods except for SpatialDE, and to a lesser extent nnSVG and Giotto KM, demonstrate a high fidelity in gene ranks across all datasets. Therefore, the methods that have a lower correlation when the genes included in a dataset changes, do not independently calculate the SVG statistics for each gene (**Figure 4b**). Although there is some variability in MERINGUE, SPARK-X, Moran’s I, Giotto rank and SOMDE, this variability may not have a significant impact on downstream analysis. These analyses reveal that the decisions made on gene filtering, a common step in data pre-processing, may result in a change in SVG statistics and their ranking for some of the SVG detection methods.

**Figure 4.**
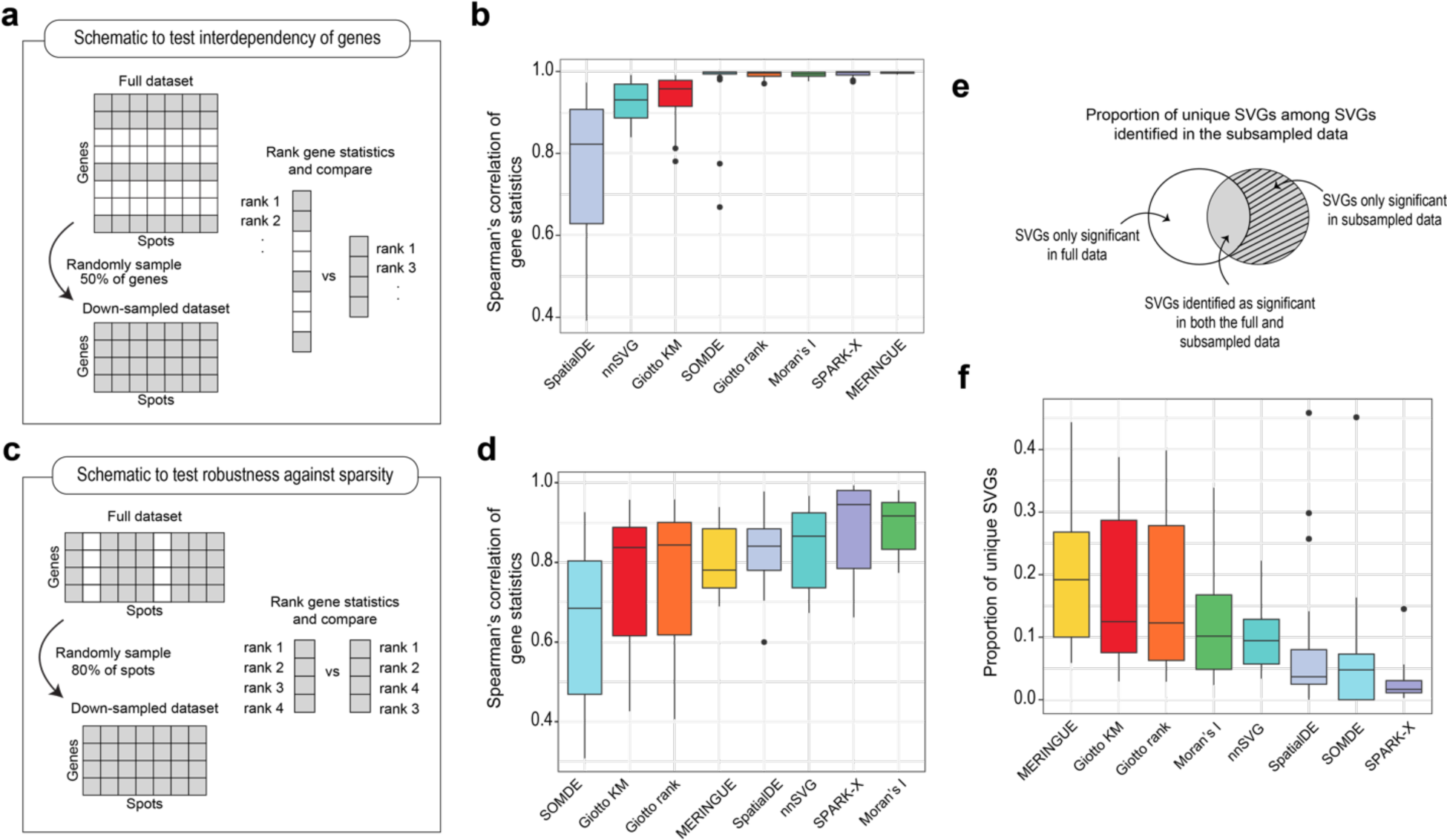
Reproducibility of SVG detection tools with down-sampling of the data. **(a)** Schematic of the experimental approach to determine the interdependency of genes in the calculation of SVG statistics. For the down-sampling, 50% of the genes were randomly down-sampled whilst keeping the number of spots equal. **(b)** Boxplots of the Spearman correlation results performed on all datasets coloured by SVG method and ordered by increasing mean correlation coefficient. **(c-d)** as in **(a-b)** but for the investigation of the reproducibility of SVG methods with increased sparsity in spatial spots. The datasets were randomly down sampled to 80% of the total number of spots. **(e)** Venn diagram illustrates the proportions of uniquely identified SVGs as described below in **(f)**. Boxplots of the proportion of false positive SVGs calculated as the proportion of SVGs uniquely identified in the down-sampled data divided by all the significant SVGs identified in the down-sampled data.

Each spatial technology has a different capacity to capture spatial locations (**Supplementary Figure 1a**) which may be due to the relatively low-throughput nature of some spatial technologies or inefficiencies in sample preparation. To test the robustness of each method against the sparsity of spatial locations, we down-sampled all datasets to 80% of the total number of spatial spots and repeated the SVG detection (**Figure 4c**).

Across all methods, there is some degree of variability in the spearman’s correlation amongst datasets due to the induced sparsity (**Figure 4d**). In particular, we found that the variability amongst datasets and the degree of sensitivity to spot sparsity tend to be greater for methods that rely on neighbourhood adjacency relationships like nnSVG (uses spatial covariance functions in Gaussian Processes using a nearest-neighbour Gaussian process model), SOMDE (uses self-organising map to cluster neighbouring cells into nodes), MERINGUE (uses neighbourhood relationships encoded by a Voronoi Tessellation and Delaunay-derived weighted adjacency matrix), and the Giotto methods (uses a Delaunay triangulation network based on cell centroid physical distances). Conversely, methods that were less sensitive were SPARK-X, SpatialDE, and Moran’s I. The reliance on such nearest neighbourhood maps or distance-based networks in the former group of methods may explain the sensitivity to sparsity as it affects the detection of SVGs based on its expression between neighbours in a spatial network.

To investigate the capacity of the methods to correctly identify SVGs and avoid the detection of false positive SVGs with induced down-sampling of the spatial spots, we next quantified the proportion of SVGs that are uniquely identified in the down-sampled data (**Figure 4e**). We consider that the original full dataset has the most power to detect SVGs and any significant SVGs that are detected in the down-sampled data but not in the original data are false positives. We visualised the proportions of all significant SVGs identified in the down-sampled data that are either identified as significant in the full data or unique to the down-sampled data (**Figure 4f**). Our findings show that SPARK-X, SOMDE and SpatialDE performed the best in terms of identifying the lowest proportion of false positive SVGs with down-sampling of the data. Although the performance of SOMDE suffers under induced sparsity, the low proportion of false positive SVGs may be explained by the fact that SOMDE tends to select fewer SVGs overall compared to other methods (**Fig 2b**). Again, for most methods there is high variability amongst datasets, which suggests that a method’s performance may be dataset dependent under sparse conditions.

Overall, our down-sampling experiments of genes and spots show that the performance of most methods to detect significant SVGs may be affected by changes in the gene number and sparsity of spatial spots. This has important implications when considering the most suitable method that is insensitive to gene filtering and dataset quality.

### Accuracy of SVG methods in detecting SVGs using synthetic spatial transcriptomics data

To test the accuracy of the SVG detection methods, we next simulated spatial transcriptomics datasets with ground truth SVGs and spatially invariant genes using scDesign3^19^ (**Supplementary Figures 6-8**). To enable representation of the diverse sequencing technologies and tissue histologies in real spatial data, we simulated in silico data from nine data sources covering nine distinct spatial masks, five tissue histology types, two spatial platform technologies, and diverse sequencing depths (590-1937 genes, 59-194 spatially variable genes, and 369-4895 spatial spots). We then performed SVG detection on the simulated datasets using the eight methods and evaluated their performance by calculating the true positive rate (TPR) and the false discovery rate (FDR) across three adjusted *p*-value thresholds (0.01, 0.05, and 0.1) (see Methods for details). At the adjusted *p*-value thresholds of 0.01 and 0.05, we found that SPARK-X, SOMDE, nnSVG, and SpatialDE performed well with a high TPR and a low FDR (**Figure 5 and Supplementary Figure 9**). Under adjusted *p*-value thresholds of 0.01, 0.05, and 0.1, Giotto rank, Moran’s I, and nnSVG all demonstrated a high TPR but suffered from a high level of false positive identification. Compared to the other methods, the Giotto methods and Moran’s I performed relatively poorly in the simulation, displaying the highest FDRs in most datasets (**Figure 5a and b**). These methods tended to identify a greater proportion and number of significant SVGs (**Supplementary Figure 9b and c**). These findings reveal that for methods, except SPARK-X and SOMDE, the estimated FDRs (i.e., adjusted *p*-value thresholds) do not accurately represent the true FDRs for SVG detection in these simulated datasets.

**Figure 5.**
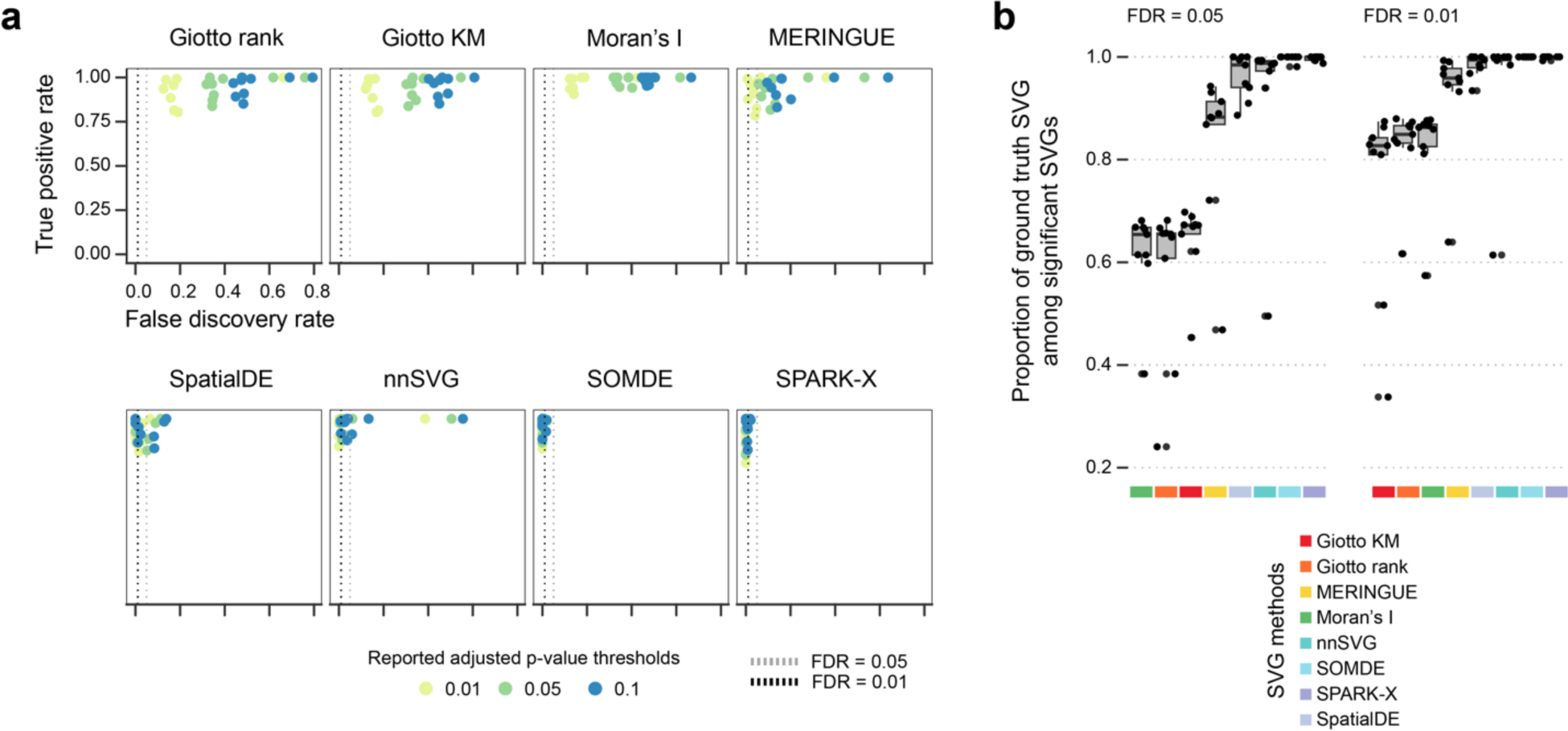
Spatially variable gene detection performance across 9 simulated datasets. **(a)** Scatter plot of observed true positive rate (y-axis) and false discovery rate (x-axis) of spatially variable gene detection by the benchmarked tools at six adjusted p-value cut-offs at 0.1, 0.01, and 0.05. Each dot is colour-coded by the cut-off used. The two horizontal lines represent the true FDRs of 0.01 and 0.05. **(b)** The proportion of ground truth SVGs among significant SVGs determined by each method at an FDR-adjusted p-value of 0.01.

### Performance on clustering spatial domains

A key task in spatial transcriptomics data analysis is to identify spatial domains that mark distinctive cell and tissue types in a biological sample. One approach to achieve this is to cluster profiled locations into spatial domains using SVGs. To compare the capacities of SVGs identified by each method in clustering the spatial domains, we took advantage of the spatial transcriptomics data of an E9.5 mouse embryo given the availability of tissue annotations in these samples (**Figure 6a**). First, we performed SVG calling using each SVG detection method. Then taking a varying number of top SVGs, we computed the top 20 principal components (PCs) using the feature-selected spatial transcriptomics data. Using either spatially-aware clustering tools (BayesSpace^20^ and SpaGCN^21^) or canonical clustering approaches (k-means, hierarchical, Louvain, and Leiden clustering using the SINFONIA framework^22^), we performed clustering on the top 20 PCs and calculated the concordance between the clustering results and the pre-defined spatial domains to measure the performance of the SVGs to delineate the anatomical locations. By taking a large range in the number of features used (between 100 and 1900 features), we were able to observe an overall increasing trend in performance with an increasing number of SVGs used for all SVG methods, with the accuracy in classification peaking at around 900-1100 SVGs (**Figure 6b**). Whilst this observation was broadly consistent, the pattern differed for some clustering and SVG method combinations. For example, hierarchical clustering demonstrated a decreasing trend in accuracy with increasing number of SVGs used unlike most clustering methods. The overall pattern was consistent between different concordance measures, including Fowlkes-Mallows index (FMI), normalised mutual information (NMI), and purity score (**Supplementary Figure 10).** These results suggest that whilst the selection of the number of top SVGs used in clustering will depend on the data using approximately between 900-1300 genes for the dataset tested led to the highest accuracy in clustering of spatial domains across most conditions.

**Figure 6.**
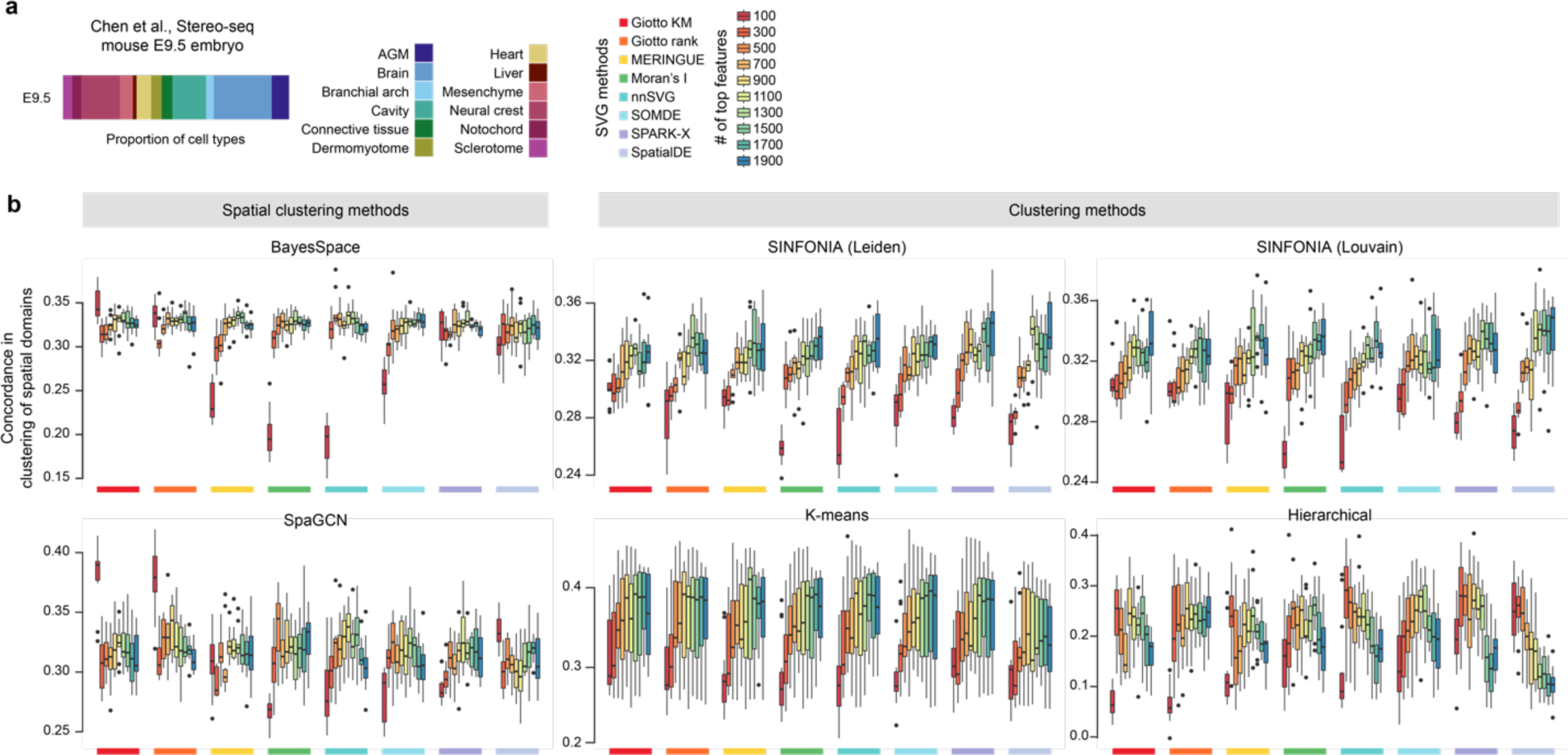
Performance of SVGs selected by each method for clustering spatial domains in mouse embryos. (**a**) Proportion spatial locations annotated to one of sixteen tissue domains in the E9.5 mouse embryo. Concordance in the clustering outputs and the pre-defined spatial domains in the mouse embryo was computed across a range of top SVGs (between 100 and 1900 genes) selected by each method. Clustering is performed using two spatial clustering methods (BayesSpace and SpaGCN) and four non-spatial clustering methods (SINFONIA’s Louvain and Leiden, k-means, and hierarchical clustering). Concordance between the clustering outputs and the pre-defined spatial domains is quantified in terms of the adjusted Rand index.

### Computational time and memory usage

Computational time and memory usage are key considerations in practical applications, especially for large spatial transcriptomics data analyses. In our evaluation, we configured a standard virtual machine, with 16 OCPUs and 256 GB of memory and recorded the runtime and the peak memory usage for each SVG detection method on each dataset (**Figure 7**). As expected, we found the computational time and the peak memory usage are both positively correlated with the number of spatial locations in the datasets. In terms of computational time, comparison across methods revealed that SPARK-X is the fastest method and scales extremely well with the number of spatial locations. While SOMDE is the second best in most cases, it is significantly slower compared to SPARK-X. In contrast, SpatialDE performed poorer especially on datasets with large numbers of spatial locations. Giotto KM performed poorly in most of the datasets but does scale better than SpatialDE with the number of spatial locations in datasets. Similarly, nnSVG scaled better with the number of spatial locations than SpatialDE but was slower on datasets with many genes. In terms of peak memory usage, we found that SOMDE uses the least peak memory across all datasets and SPARK-X ranked the second in most cases although it has a significantly higher peak memory usage. In comparison, the two methods implemented in Giotto and SpatialDE show high peak memory usage especially in datasets with many spatial locations. While there is a trade-off between speed and memory usage, taken together, these results suggest that SPARK-X and SOMDE are the most efficient methods in terms of speed and memory usage for SVG detection.

**Figure 7.**
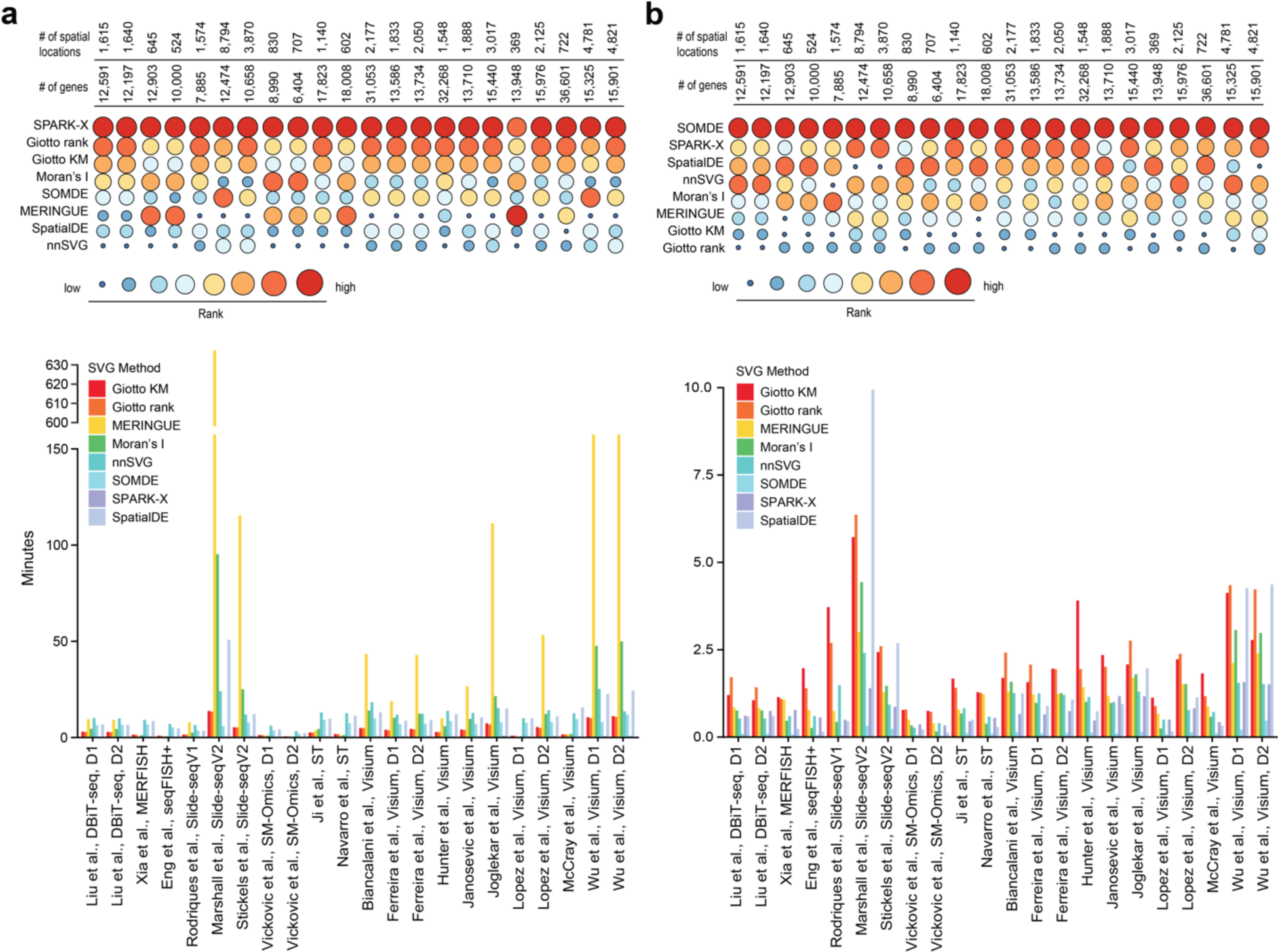
Evaluation of computational speed and peak memory usage of SVG detection methods. (**a**) Computation time in units of minutes and (**b**) peak memory usage in units of GiB across datasets.

## Discussion

We found that, for most methods, a significant proportion of genes were detected as SVGs under the adjusted *p*-value of 0.05 in most of the tested datasets (**Figure 2b** and **Supplementary Figure 3c**). However, the overlaps across the eight methods were relatively small considering the large numbers of SVGs identified from each SVG detection method (**Figure 2c**), suggesting large discrepancies among SVG detection methods when a significance cutoff is used to filter for SVGs. Consistent with this, in our simulation study where ground truth SVGs were introduced into simulated spatial transcriptomics data, we found that for some methods, in particular the Giotto methods and Moran’s I, the estimated FDRs did not accurately represent the true FDRs in most of the synthetic datasets (**Figure 5**). These results highlight that the estimation of statistical significance is difficult and there is much room for improvement. It also cautions the use of and the reliance on such statistical significance from some of the current SVG detection tools for drawing data and biological conclusions.

We also discovered that SVGs identified by most methods show a strong positive correlation with their expression levels (**Figure 3**). We note that a similar relationship was found between gene variability and expression level in scRNA-seq data and most computational methods designed for HVG detection actively correct for such a ‘bias’ ^2^. While we could not rule out the possibility that genes that vary spatially are also highly expressed, future work should be performed to investigate the biological basis and plausibility for such a correlation. During practical application, it is important to be aware of the tendency of current SVG detection tools to select genes with high expression levels. Future method development will be required to account for this effect such as to retain relatively lowly expressed genes such as transcription factors in downstream analysis. In addition, we found that for most methods the relative rankings of SVGs change when different pools of genes and spots are included in the datasets (**Figure 4c and f**). While considering the interdependency among genes may provide useful information for identifying SVGs, it is important to be aware that different SVG detection results may be obtained when different preprocessing steps were used to filter genes prior to SVG analysis.

Lastly, SVG detection can be viewed as a feature selection step in spatial transcriptomics data analysis, where useful features (i.e., SVGs) are selected and/or uninformative ones are removed. In particular, the current SVG detection methods can be considered as unsupervised approaches where no information such as cell types, cell states, or spatial domains are required. A great amount of work has been done in feature selection in single-cell data analysis ^23^, including unsupervised methods and also more advanced methods that perform combinatorial feature selection using supervised learning such as embedded feature selection using random forest and wrapper feature selection using genetic algorithms. We anticipate that future development of SVG detection methods will explore the utility of information such as cell types and states to identify SVGs that not only independently mark the spatial variability but also those that cooperate across multiple genes and together define spatial variability. We believe these developments will introduce additional computational new challenges but will undoubtedly lead to new biological insight from spatial transcriptomics data analyses.

## Methods

### Datasets

Summary information of spatial transcriptomics datasets was included in **Supplementary Figure 1a**. Below we provide the accession numbers when available or download links used to obtain each dataset.

- Liu et al., DBiT-seq^18^. Mouse Embryo E12 (GSM4189614_0628cL) and E11(GSM4364243_E11-2L). Downloaded from GEO accession GSE137986.
- *Xia et al., MERFISH* ^14^. Human osteosarcoma. Downloaded from the supplementary section of the corresponding paper. https://www.pnas.org/doi/suppl/10.1073/pnas.1912459116/suppl_file/pnas.1912459116.sd12.csv
- *Eng et al., SeqFISH+* ^15^. Mouse primary visual cortex (VISp). Downloaded from https://github.com/CaiGroup/seqFISH-PLUS. The spatial coordinate of each spot was generated using ‘stitchFieldCoordinates’ function in Giotto.
- Rodriques et al., SlideseqV1^12^. Mouse cerebellum. Downloaded the ‘Puck_180819_11’ sample from https://singlecell.broadinstitute.org/single_cell/study/SCP354/slide-seq-study.
- Marshall et al., SlideseqV2^13^. Human kidney cortex. Downloaded the ‘HumanKidney_Puck_20011308’ sample from https://cellxgene.cziscience.com/datasets.
- *Stickels et al., Slide-seqV2* ^24^. Mouse hippocampus. Downloaded the ‘Puck_200115_08’ sample from https://singlecell.broadinstitute.org/single_cell/study/SCP815/highly-sensitive-spatial-transcriptomics-at-near-cellular-resolution-with-slide-seqv2#study-download
- Vickovic et al., SM-Omics^17^. Mouse brain cortex. Downloaded the ‘10015CN78_C1_stdata_adjusted’ and ‘10015CN89_D2_stdata_adjusted’ samples from https://singlecell.broadinstitute.org/single_cell/study/SCP979/sm-omics-an-automated-platform-for-high-throughput-spatial-multi-omics.
- *Ji et al., ST* ^11^. Human squamous carcinoma. Downloaded from GSM4284322.
- *Navarro et al., ST* ^25^. Mouse hippocampus wild-type replicate 1. Downloaded from https://data.mendeley.com/datasets/6s959w2zyr/1.
- *Biancalani et al., Visium* ^26^. Mouse primary motor cortex. Downloaded from https://storage.googleapis.com/tommaso-brain-data/tangram_demo/Allen-Visium_Allen1_cell_count.h5ad
- *Ferreira et al., Visium* ^27^. Mouse kidney. Downloaded the Sham model and ischemia reperfusion injury model from GSE171406.
- *Hunter et al., Visium* ^28^. Zebrafish melanoma. Downloaded the ‘Visium-A’ sample from GSE159709.
- *Janosevic et al., Visium* ^29^. Mouse kidney. Downloaded from GSE154107.
- *Joglekar et al., Visium* ^30^. Mouse pre-frontal cortex. Downloaded from GSE158450
- *Lopez et al., Visium* ^31^. Mouse lymph node and MCA205 tumour. Downloaded from GSE173776 and GSE173773 respectively.
- *McCray et al., Visium* ^32^. Human prostate. Downloaded from GSM4837767.
- *Wu et al., Visium* ^33^. Human breast cancer. https://zenodo.org/record/4739739#.YY6N_pMzaWC
- *E9.5 Mouse Embryo* ^16^. E9.5 mouse embryo spatial profile. Downloaded from https://db.cngb.org/stomics/mosta/.

### SVG detection methods

Datasets were first filtered by first removing cells whose top-50 highly expressed genes contributed to 50% of the total counts and then genes that were expressed in fewer than 30 cells. Log normalisation of raw counts was performed prior to SVG detection as per the recommended default for each method. The same reproducible seed was set prior to running each method.

#### Giotto KM and Giotto rank

Giotto^7^ requires a spatial Delaunay triangulation network to be built on reduced dimensions to represent the spatial relationships. Then, statistical enrichment using Fisher’s exact test of binarized expression in spatial nearest neighbours is performed to determine SVGs. The two methods differ in their binarization method. In *Giotto KM*, expression values for each gene are binarized using k-means clustering (k=2), otherwise simple thresholding on rank is applied in *Giotto rank* (default = 30%). Thus, a gene is considered an SVG if it is highly expressed in neighbouring cells. Normalisation was performed using *normalizeGiotto()* under default parameters. SVG detection was thus performed with two different approaches k-means and rank using *binSpect(bin_method=”kmeans”)* and *binSpect(bin_method=”rank”)* respectively, following the author’s tutorial. https://rubd.github.io/Giotto_site/articles/mouse_visium_kidney_200916.html

### Moran’s I

Moran’s I ranks genes by the observed spatial autocorrelation ^34,35^ to measure the dependence of a feature on spatial location. Weights are calculated as 1/distance. Raw counts were first normalised using *SCTransform()*. Using Seurat v4.1.1, SVGs were detected using *FindSpatiallyVariableFeatures(selection.method = “moransi”)* and statistics for all features were returned. p-value adjustment was manually performed using the BH method.

#### MERINGUE

MERINGUE identifies spatially variable genes using neighbourhood adjacency relationships and spatial auto-correlation. MERINGUE first represents cells as neighbourhoods using Voronoi tessellation. Then, the resulting Delaunay-derived weighted adjacency matrix and a matrix of normalised gene expression is used to calculate Moran’s I. Raw counts were CPM-normalised using *scuttle::normalizeCounts()* and the default filtering distance was used to generate the weighted adjacency matrix. Statistics and p-values for all features were returned. P-value adjustment was manually performed using the BH method.

#### nnSVG

nnSVG is based on scalable estimation of spatial covariant functions in Gaussian process regression using nearest neighbour Gaussian process (NNGP) models. The BRISC algorithm^36^ was used to implement the NNGP model and obtain maximum likelihood parameter estimates for each gene. A likelihood-ratio test is performed to rank genes by estimated LR statistic values. Log normalisation was performed using *scater::LogNormCounts* prior to running *nnSVG()* with default parameters (k=10). Where default parameters were unsuccessful, the number of nearest neighbours was fine-tuned from k=5 to k=15.

#### SOMDE

A SOM neural network is used to adaptively integrate nearest neighbour data into different nodes, achieving a condensed representation of the spatial transcriptome. SVGs are identified on a node-level, using spatial location and gene meta-expression information. A squared exponential Gaussian kernel is applied to generate log-likelihood ratio values wherein a likelihood-ratio test is performed to rank genes by estimated LLR statistic values. The procedure was performed as per the recommended tutorial at https://github.com/WhirlFirst/somde using python. k=10 was chosen as the default nearest neighbours when constructing the SOM across all benchmarking datasets to preserve local spatial patterns across both small and large datasets. Where default parameters were unsuccessful, the number of nearest neighbours was fine-tuned from k=5 to k=20.

#### SPARK-X

SPARK-X is a non-parametric method that relies on a robust covariance test framework, including the Hilber-Schmidt independence criteria test and the distance covariance matrix test. A test statistic is observed by measuring the similarity between two relationship matrices based on gene expression and spatial coordinates respectively. A p-value is computed for each distance covariance matrix constructed and a Cauchy combined p-value is reported. *sparkx()* was run under default parameters.

#### SpatialDE

SpatialDE fits a linear mixed model for each gene with gaussian kernels and decomposes the gene variation into spatial or non-spatial variation. The non-spatial variation is separately modeled using observed noise, and the spatial variation is explained by an exponential covariance function. For each gaussian kernel, a p-value is calculated from the likelihood test to rank genes by estimated LR statistics. SpatialDE was run under the python implementation and the procedure was as follows in the tutorial by the authors as in https://github.com/Teichlab/SpatialDE.

### Correlation of ranked gene statistics

To calculate the pairwise spearman’s correlation between each method for each dataset, the corresponding gene statistics were used as outlined in Table 1. Where a comparable gene statistic was not reported by a method, the -log10(adjusted p-value) was used to rank the genes.

**Table 1.** Description of statistics used to rank genes and the p-value adjustment methods used by each package.

| SVG Method | Statistic | p-value adj. |
| --- | --- | --- |
| Giotto k-means | $-\log_{10}(\text{adjusted p-value})$ | BH |
| Giotto rank | $-\log_{10}(\text{adjusted p-value})$ | BH |
| MERINGUE | observed coefficient | BH |
| Moran's I | observed coefficient | BH |
| nnSVG | LLR statistic | BH |
| SOMDE | LLR statistic | q-value |
| SPARK-X | $-\log_{10}(\text{adjusted p-value})$ | BY |
| SpatialDE | LLR statistic | q-value |

### Identifying significant SVGs

Significant SVGs were typically defined as genes with an adjusted p-value of < 0.05. Specifically for Moran’s I, genes that have a positive spatial autocorrelation coefficient and an adjusted p-value of < 0.05 were selected as significant.

### Dependency across genes

To assess the dependency across genes in SVG analysis, we randomly down-sampled 50% of genes from all datasets that ran successfully. We next applied each SVG detection method and calculated SVG statistics of remaining genes in the down-sampled dataset as per Table 1. The relative rank of these genes was compared with their rank in the original full dataset to assess if there is any change of relative ranking when other genes were included in the dataset. Methods that lead to a different ranking of SVGs in the down-sampled dataset when additional genes were included are considered as calculating spatial variability of a gene depending on the presence and absence of other genes.

### Robustness against sparsity

To assess how each method performs against sparse data, we randomly down-sampled 80% of spots from all datasets that ran successfully. After applying each SVG detection method, we evaluated the performance of each method in two aspects. To assess the impact of sparsity on the relative rankings of the gene statistics, we computed the spearman’s correlation of the original dataset and the down-sampled dataset using the statistics reported in Table 1. To assess the extent of sparsity on the significantly detected SVGs, we visualised the proportion of uniquely detected SVGs because of the subsampling against the total number of SVGs significantly detected in the original dataset.

### Simulation of spatial transcriptomics data

To evaluate the capacity of methods to detect SVGs with high sensitivity and specificity, we simulated a set of spatial transcriptomics data using scDesign3^19^, providing us with ground truth spatially variable genes. The synthetic data were generated using real spatial transcriptomics datasets from **Supplementary Figure 1a**. Simulation of realistic spatial transcriptomics data was performed following the default settings of scDesign3. To enable fast computation of the model parameters estimated from the real data, we simulated up to approximately 2000 genes and for each dataset generated 10% of all genes as spatially variable. The synthetic datasets model parameters from nine datasets from seven independent studies that cover different sequencing technologies (Visium and DBiT-seq), tissue histologies (breast cancer, brain, embryo, and cancer), number of spatial spots (369-4895 spatial spots) and sequencing depths (590-1937 genes and 59-194 spatially variable genes).

### Benchmarking of simulation studies

To evaluate the performance of the SVG detection methods on the simulated data, we calculated the receiver operating characteristic curve based on the statistics or *p*-values of the genes, indicating the capacity of methods to rank truly spatially variable genes before non-variable ones. We next calculated the true positive rate (TPR) and the false discovery rate (FDR) to evaluate FDR control at six adjusted *p*-value thresholds (1e-100, 1e-50, 1e-10, 0.01, 0.05, and 0.1) for each simulated dataset. The cutpointr package ^37^ was used to calculate the TPR and FDR performance metrics.

### Clustering and concordance quantification

To quantify the utility of SVGs in spatial domain clustering, we used varying number of top significant SVGs (between 100 and 1900 genes) reported from each method to subset the expression matrix, compute principal component analysis, and performed clustering on the top 20 principal components to cluster the E9.5 mouse embryo spatial transcriptomics data into 13 tissue domains based on the original annotation^16^. We performed 10 repeats by random subsampling of the spatial data to 80% of the total number of spatial spots for each repeat. We performed either spatial clustering using the default settings (unless otherwise stated) of BayesSpace^20^ (gamma = 2 and nrep = 1000) and SpaGCN^21^ or *k*-means, hierarchical, Louvain, and Leiden clustering. The total number of clusters was set to the total number of spatial domains observed in the data. In particular, we performed a binary search to tune the resolution parameter as described in SINFONIA^22^ to tune the clustering in the two community-based clustering algorithms. To assess the clustering performance of the SVGs defined by various SVG detection methods, we used the adjusted Rand index (ARI), the normalized mutual information (NMI), the Fowlkes-Mallows index (FMI), and purity to evaluate the concordance between the clustering labels and the spatial domains. Each metric was calculated as follows:

#### Adjusted Rand index

Let *T* denote the known ground-truth spatial domains of spots, *P* denote the predicted clustering labels from k-means clustering, *N* denote the total number of spatial locations, *x_i_* denote the number of spots assigned to the *i*th cluster of *P*, *y_i_* denote the number of spots that belong to the *j*th unique label of *T*, and *n_ij_* denote the number of overlapping spots between the *i*th cluster and the *j*th unique label. The Rand index (RI) denotes the probability that the obtained clusters and the spatial domain labels agree on a randomly chosen pair of spots. The adjusted Rand index (ARI) adjusts for the expected agreement by chance.

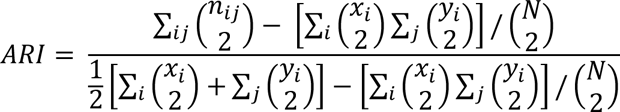

#### Normalised mutual information

Normalised mutual information (NMI) assesses the similarity between the obtained cluster labels and the ground-truth spatial locations, scaled between 0 and 1. We calculate the NMI as follows:

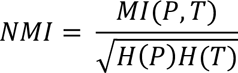

Where *H*(.) is the entropy function.

A comparison of ARI and NMI presented in previous studies^38,39^ suggest ARI is preferred when there are large equal-sized clusters, whilst NMI is preferred in the presence of class imbalance and rare clusters.

#### Fowlkes-Mallows index

The Fowlkes-Mallows index (FMI) measures the similarity in two clustering results and is defined as the geometric mean of the precision and recall. The FMI is calculated using the following equation:

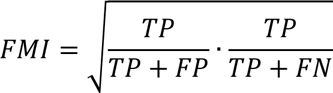

Where TP is the number of true positives, which are pairs of spots that are in the same spatial domain in both the true and predicted labels; FP is the number of false positive, which are pairs of spots that are in the same cluster in the predicted clusters but in different clusters in the ground-truth labels; and FN is the number of false negatives, which are pairs of spots that are in the same cluster in the ground-truth labels but in different clusters in the predicted clusters. The score is adjusted to a range between 0 to 1, where a value of 1 signifies when all the spatial spots are correctly labelled. A higher FMI denotes a greater similarity between the two clustering results.

#### Purity

Purity is scored in terms of whether the clusters contain only spots of the same spatial domain. Purity equals to 1 if all the spots within the same cluster correspond to the same spatial domain. The purity score is computed using the following equation:

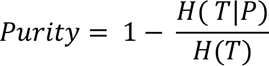

Where *H*(*T*|*P*) indicates the uncertainty of true labels based on the predicted labels.

### Time consumption and memory usage

To measure computational consumption for each method, a standard virtual machine with 16 OCPUs and 256 GB was used. Where methods offered parallelization (Giotto, SPARK-X, nnSVG, SOMDE, and SpatialDE) all available cores when it was possible to specify, were utilised to record the running time. For all methods run in R, the elapsed time to run each method was evaluated using the *system.time()* function. The peak memory usage was monitored using *gc()*. For methods run in python, *perf_counter()* from the *time* package was used to record the elapsed time. To record the peak memory usage, *get_traced_memory()* was used from the *tracemalloc* package.

## Acknowledgment

We thank our colleagues at the School of Mathematics and Statistics, The University of Sydney, and the Sydney Precision Bioinformatics Alliance for their feedback. This work is supported by a National Health and Medical Research Council (NHMRC) Investigator Grant (1173469) and a Metcalf Prize to P.Y.; and a postgraduate scholarship from Research Training Program and a Children’s Medical Research Institute postgraduate scholarship to C.C.

## Author’s contribution

C.C., H.J.K., and P.Y. conceived the study. C.C. and H.J.K. performed data analyses and interpreted the results with input from P.Y. All authors wrote the manuscript and approved the final version of the manuscript.

## References

1. Svensson, V., Teichmann, S. A. & Stegle, O. SpatialDE: identification of spatially variable genes. Nat Methods 15, 343–346 (2018).

2. Yip, S. H., Sham, P. C. & Wang, J. Evaluation of tools for highly variable gene discovery from single-cell RNA-seq data. Brief Bioinform 20, 1583–1589 (2019).

3. Kolodziejczyk, A. A., Kim, J. K., Svensson, V., Marioni, J. C. & Teichmann, S. A. The Technology and Biology of Single-Cell RNA Sequencing. Molecular Cell 58, 610–620 (2015).

4. Sun, S., Zhu, J. & Zhou, X. Statistical analysis of spatial expression patterns for spatially resolved transcriptomic studies. Nat Methods 17, 193–200 (2020).

5. Zhu, J., Sun, S. & Zhou, X. SPARK-X: non-parametric modeling enables scalable and robust detection of spatial expression patterns for large spatial transcriptomic studies. Genome Biol 22, 184 (2021).

6. Hao, M., Hua, K. & Zhang, X. SOMDE: A scalable method for identifying spatially variable genes with self-organizing map. Bioinformatics btab471 (2021) doi:10.1093/bioinformatics/btab471.

7. Dries, R. et al. Giotto: a toolbox for integrative analysis and visualization of spatial expression data. Genome Biol 22, 78 (2021).

8. Weber, L. M., Saha, A., Datta, A., Hansen, K. D. & Hicks, S. C. nnSVG: scalable identification of spatially variable genes using nearest-neighbor Gaussian processes. http://biorxiv.org/lookup/doi/10.1101/2022.05.16.492124(2022) doi:10.1101/2022.05.16.492124.

9. Miller, B. F., Bambah-Mukku, D., Dulac, C., Zhuang, X. & Fan, J. Characterizing spatial gene expression heterogeneity in spatially resolved single-cell transcriptomics data with nonuniform cellular densities. Genome Res. gr.271288.120 (2021) doi:10.1101/gr.271288.120.

10. Hao, Y. et al. Integrated analysis of multimodal single-cell data. Cell 184, 3573–3587.e29 (2021).

11. Ji, A. L. et al. Multimodal Analysis of Composition and Spatial Architecture in Human Squamous Cell Carcinoma. Cell 182, 497–514.e22 (2020).

12. Rodriques, S. G. et al. Slide-seq: A scalable technology for measuring genome-wide expression at high spatial resolution. Science 363, 1463–1467 (2019).

13. Marshall, J. L., et al. High-resolution Slide-seqV2 spatial transcriptomics enables discovery of disease-specific cell neighborhoods and pathways. iScience 25, (2022).

14. Xia, C., Fan, J., Emanuel, G., Hao, J. & Zhuang, X. Spatial transcriptome profiling by MERFISH reveals subcellular RNA compartmentalization and cell cycle-dependent gene expression. Proc. Natl. Acad. Sci. U.S.A. 116, 19490–19499 (2019).

15. Eng, C.-H. L. et al. Transcriptome-scale super-resolved imaging in tissues by RNA seqFISH+. Nature 568, 235–239 (2019).

16. Chen, A. et al. Spatiotemporal transcriptomic atlas of mouse organogenesis using DNA nanoball-patterned arrays. Cell 185, 1777–1792.e21 (2022).

17. Vickovic, S. et al. SM-Omics is an automated platform for high-throughput spatial multi-omics. Nat Commun 13, 795 (2022).

18. Liu, Y. et al. High-Spatial-Resolution Multi-Omics Sequencing via Deterministic Barcoding in Tissue. Cell 183, 1665–1681.e18 (2020).

19. Song, D. et al. scDesign3 generates realistic in silico data for multimodal single-cell and spatial omics. Nat Biotechnol 1–6 (2023) doi:10.1038/s41587-023-01772-1.

20. Zhao, E. et al. Spatial transcriptomics at subspot resolution with BayesSpace. Nat Biotechnol 39, 1375–1384 (2021).

21. Hu, J. et al. SpaGCN: Integrating gene expression, spatial location and histology to identify spatial domains and spatially variable genes by graph convolutional network. Nat Methods 18, 1342–1351 (2021).

22. Jiang, R., Li, Z., Jia, Y., Li, S. & Chen, S. SINFONIA: Scalable Identification of Spatially Variable Genes for Deciphering Spatial Domains. Cells 12, 604 (2023).

23. Yang, P., Huang, H. & Liu, C. Feature selection revisited in the single-cell era. Genome Biol 22, 321 (2021).

24. Stickels, R. R. et al. Highly sensitive spatial transcriptomics at near-cellular resolution with Slide-seqV2. Nat Biotechnol 39, 313–319 (2021).

25. Navarro, J. F. et al. Spatial Transcriptomics Reveals Genes Associated with Dysregulated Mitochondrial Functions and Stress Signaling in Alzheimer Disease. iScience 23, 101556 (2020).

26. Biancalani, T. et al. Deep learning and alignment of spatially resolved single-cell transcriptomes with Tangram. Nat Methods 18, 1352–1362 (2021).

27. Ferreira, R. M. et al. Integration of spatial and single-cell transcriptomics localizes epithelial cell–immune cross-talk in kidney injury. JCI Insight 6, e147703 (2021).

28. Hunter, M. V., Moncada, R., Weiss, J. M., Yanai, I. & White, R. M. Spatially resolved transcriptomics reveals the architecture of the tumor-microenvironment interface. Nat Commun 12, 6278 (2021).

29. Janosevic, D. et al. The orchestrated cellular and molecular responses of the kidney to endotoxin define a precise sepsis timeline. eLife 10, e62270 (2021).

30. Joglekar, A. et al. A spatially resolved brain region- and cell type-specific isoform atlas of the postnatal mouse brain. Nat Commun 12, 463 (2021).

31. Lopez, R. et al. DestVI identifies continuums of cell types in spatial transcriptomics data. Nat Biotechnol 40, 1360–1369 (2022).

32. McCray, T. et al. Vitamin D sufficiency enhances differentiation of patient-derived prostate epithelial organoids. iScience 24, 101974 (2021).

33. Wu, S. Z. et al. A single-cell and spatially resolved atlas of human breast cancers. Nat Genet 53, 1334–1347 (2021).

34. Moran, P. A. P. NOTES ON CONTINUOUS STOCHASTIC PHENOMENA. Biometrika 37, 17–23 (1950).

35. Gittleman, J. L. & Kot, M. Adaptation: Statistics and a Null Model for Estimating Phylogenetic Effects. Systematic Zoology 39, 227 (1990).

36. Saha, A. & Datta, A. BRISC: bootstrap for rapid inference on spatial covariances. Stat 7, e184 (2018).

37. Thiele, C. & Hirschfeld, G. cutpointr : Improved Estimation and Validation of Optimal Cutpoints in *R*. J. Stat. Soft. 98, (2021).

38. Romano, S., Vinh, N. X., Bailey, J. & Verspoor, K. Adjusting for Chance Clustering Comparison Measures. Journal of Machine Learning Research 17, 1–32 (2016).

39. Shengquan, C., Boheng, Z., Xiaoyang, C., Xuegong, Z. & Rui, J. stPlus: a reference-based method for the accurate enhancement of spatial transcriptomics. Bioinformatics 37, i299– i307 (2021).

