## Supplementary Figures for "Evaluating spatially variable gene detection methods for spatial transcriptomics data"

**a**

| Reference and platform | Species | Tissue | # Location | # Gene | Source |
|---|---|---|---|---|---|
| Liu et al., DBiT-seq, D1 | Mouse | Embryo E12 | 1,615 | 12,591 | GSE137986 |
| Liu et al., DBiT-seq, D2 | Mouse | Embryo E11 | 1,640 | 12,197 | GSE137986 |
| Xia et al., MERFISH | Human | Osteosarcoma | 645 | 12,903 | Supplementary file |
| Eng et al., seqFISH+ | Mouse | Brain cortex | 524 | 10,000 | Zenodo#2669683 |
| Rodriques et al., Slide-seqV1 | Mouse | Cerebellum | 1,574 | 7,885 | Broad Institute |
| Marshall et al., Slide-seqV2 | Human | Kidney cortex | 8,794 | 12,474 | CELLxGENE Discover |
| Stickels et al., Slide-seqV2 | Mouse | Hippocampus | 3,870 | 10,658 | Broad Institute |
| Vickovic et al., SM-Omics, D1 | Mouse | Brain cortex | 830 | 8,990 | Broad Institute |
| Vickovic et al., SM-Omics, D2 | Mouse | Brain cortex | 707 | 6,404 | Broad Institute |
| Ji et al., ST | Human | Squamous carcinoma | 1,140 | 17,823 | GSM4284322 |
| Navarro et al., ST | Mouse | Hippocampus | 602 | 18,008 | Mendeley#6s959w2zyr.1 |
| Biancalani et al., Visium | Mouse | Primary motor cortex | 2,177 | 31,053 | Google Cloud |
| Ferreira et al., Visium, D1 | Mouse | Kidney sham | 1,833 | 13,586 | GSE171406 |
| Ferreira et al., Visium, D2 | Mouse | Kidney ischemia | 2,050 | 13,734 | GSE171406 |
| Hunter et al., Visium | Zebrafish | Melanoma | 1,548 | 32,268 | GSE159709 |
| Janosevic et al., Visium | Mouse | Kidney sepsis | 1,888 | 13,710 | GSE154107 |
| Joglekar et al., Visium | Mouse | Prefrontal cortex | 3,017 | 15,440 | GSE158450 |
| Lopez et al., Visium, D1 | Mouse | Lymph node | 369 | 13,948 | GSE173776 |
| Lopez et al., Visium, D2 | Mouse | MCA205 tumour | 2,125 | 15,976 | GSE173773 |
| McCray et al., Visium | Human | Prostate | 722 | 36,601 | GSM4837767 |
| Wu et al., Visium, D1 | Human | Breast cancer | 4,781 | 15,325 | Zenodo#4739739 |
| Wu et al., Visium, D2 | Human | Breast cancer | 4,821 | 15,901 | Zenodo#4739739 |

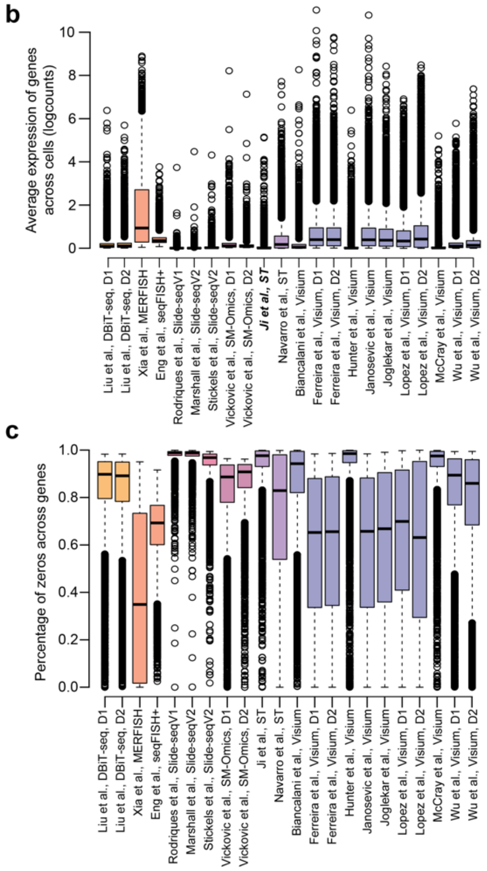


**Supplementary Figure 1. Summary information and statistics of the spatial transcriptomics datasets used for evaluating concordance, statistical significance, and reproducibility of SVG detection methods in this study.** (**a**) Dataset reference table, summarising basic information and source of each dataset. information. (**b, c**) Boxplot of mean gene expression and proportion of zero of gene across cells in each dataset.

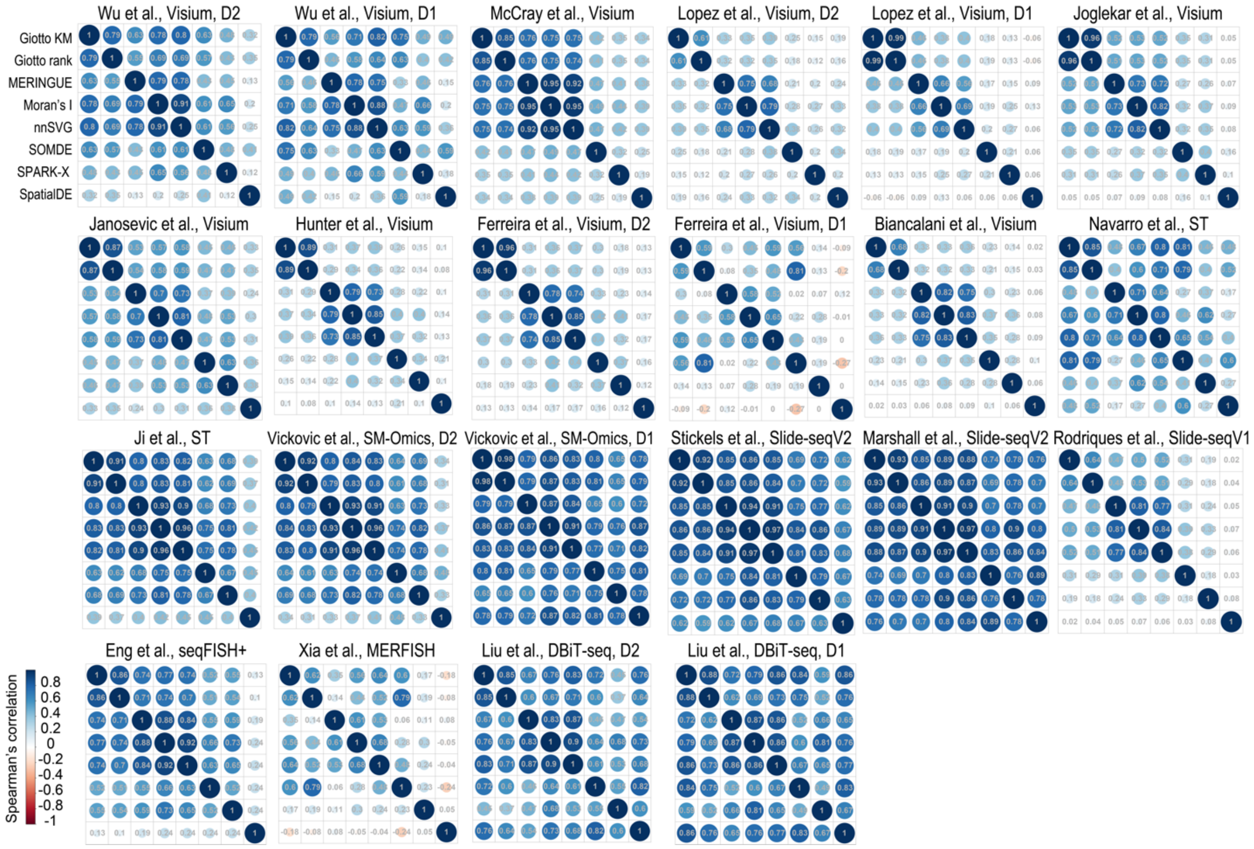


**Supplementary Figure 2. Pairwise correlation of SVG rankings reported by each method for individual spatial transcriptomics datasets.**

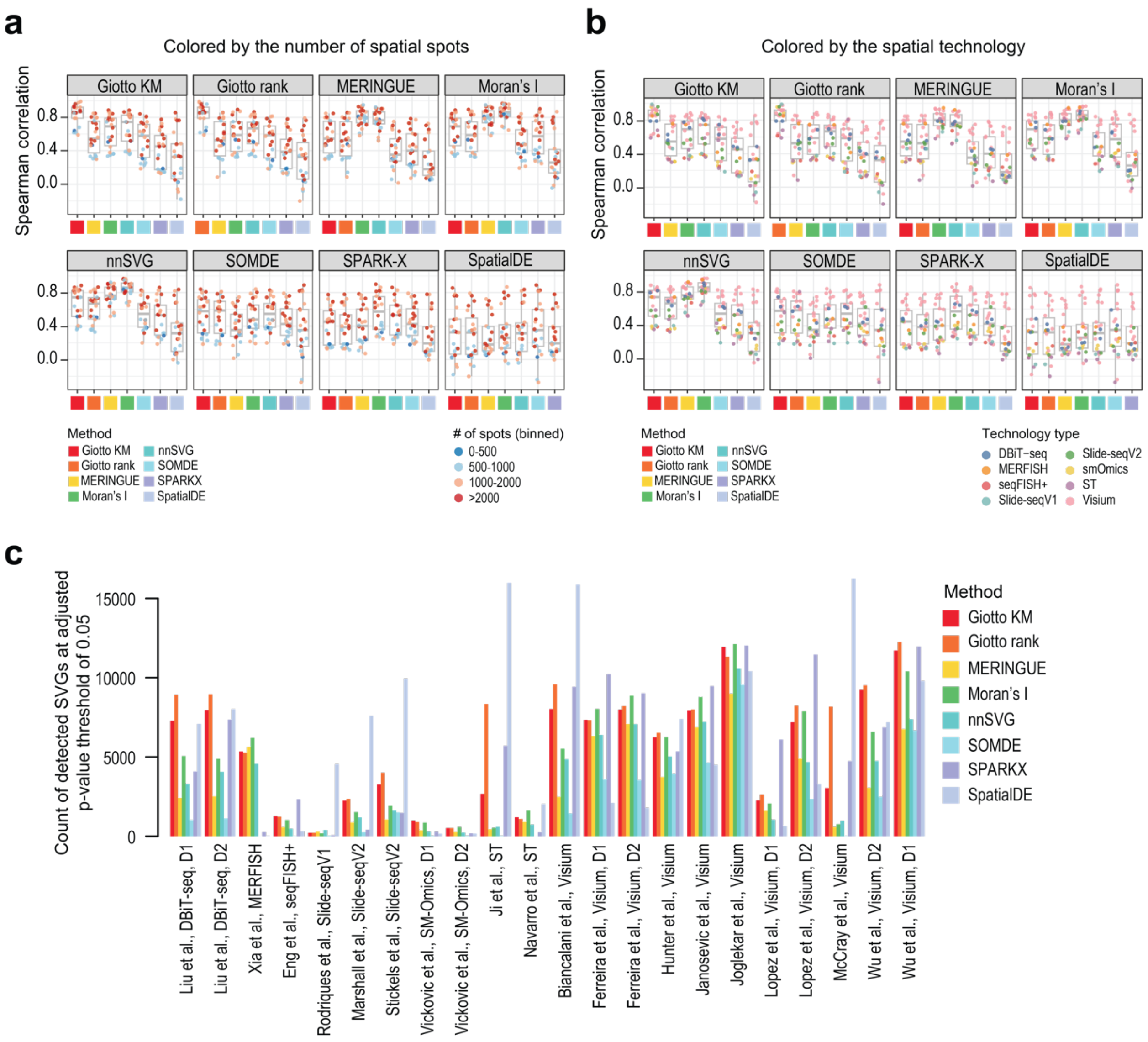


**Supplementary Figure 3. (a-b)** Boxplot of correlations of SVG rankings reported by each method against all other methods. Each dot denotes the results for a spatial transcriptomics dataset. The dots are coloured (a) by the total number of spatial spots in the dataset or (b) by the spatial technology platform. **(c)** Bar plot denoting the number of statistically significant SVGs reported by each method for each spatial transcriptomics dataset. An adjusted p-value threshold of 0.05 reported by each method for each dataset.

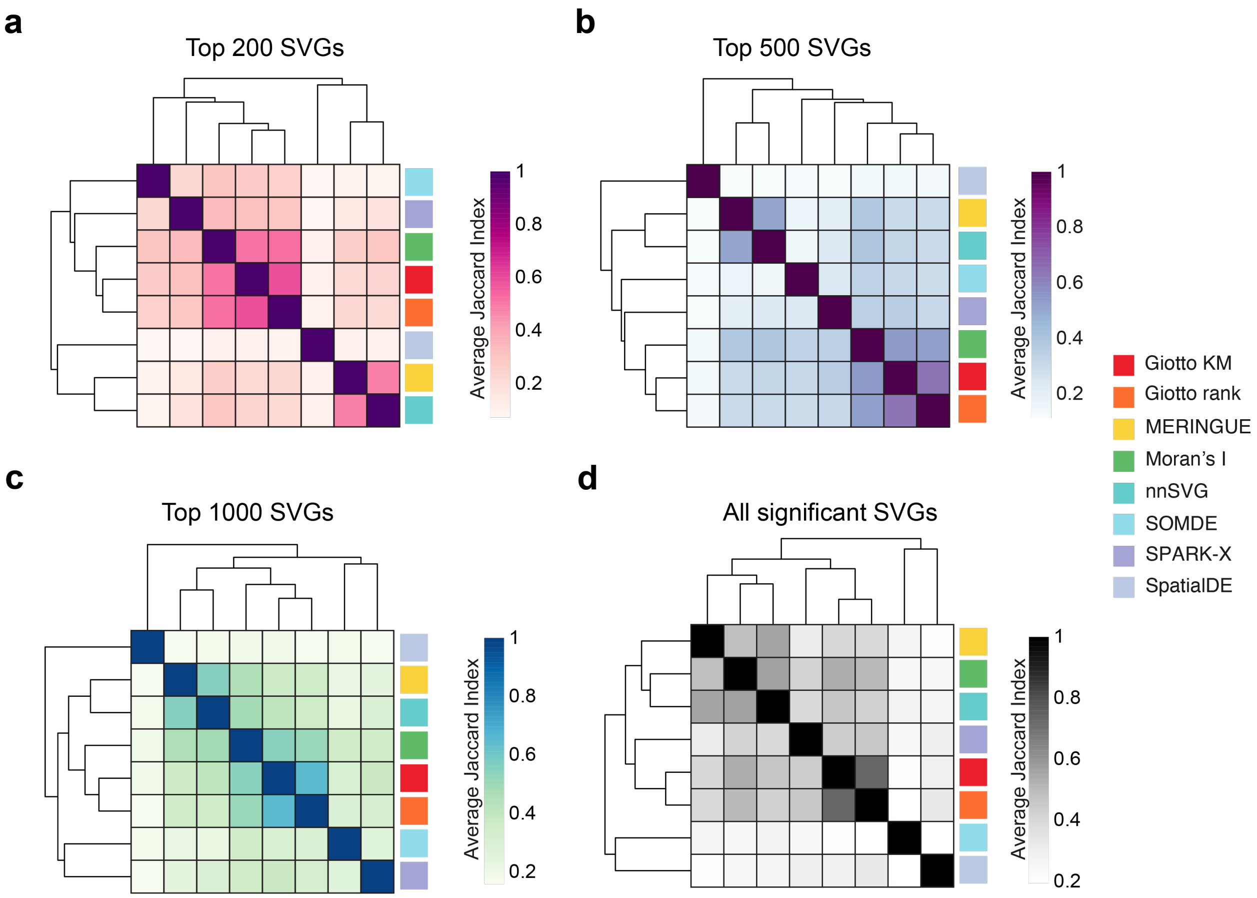


**Supplementary Figure 4. Heatmaps of the overlap of SVGs reported by each method for each spatial transcriptomics dataset.** The overlap calculated as the Jaccard Index was computed for the **(a)** top 200 SVGs, **(b)** top 500 SVGs, **(c)** top 1000 SVGs, and **(d)** all the significant SVGs (adjusted p-value <= 0.05) reported by each method for each spatial transcriptomics dataset. The average Jaccard Index across dataset and then plotted as a clustered hierarchical heatmap, where a higher Jaccard Index (darker colour) denotes a greater overlap of SVGs between methods and a lower Jaccard Index (lighter colour) denotes a lower overlap. The discrete colour bars denote the different SVG methods.

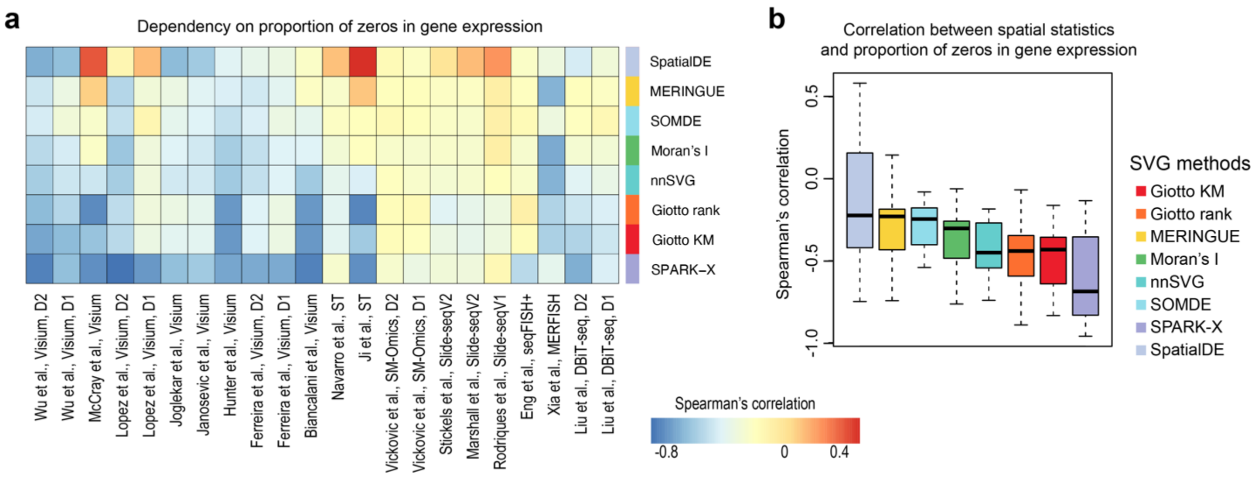


**Supplementary Figure 5. Relationship between SVG statistics and proportion of zero of genes.** (**a**) Heatmap summarising the Spearman's correlation of SVG statistics reported by each method and the proportion of zero of genes across cells in each dataset. (**b**) Boxplot of Spearman's correlation of SVG statistics reported by each method and the proportion of zero of genes across the spatial transcriptomics datasets.

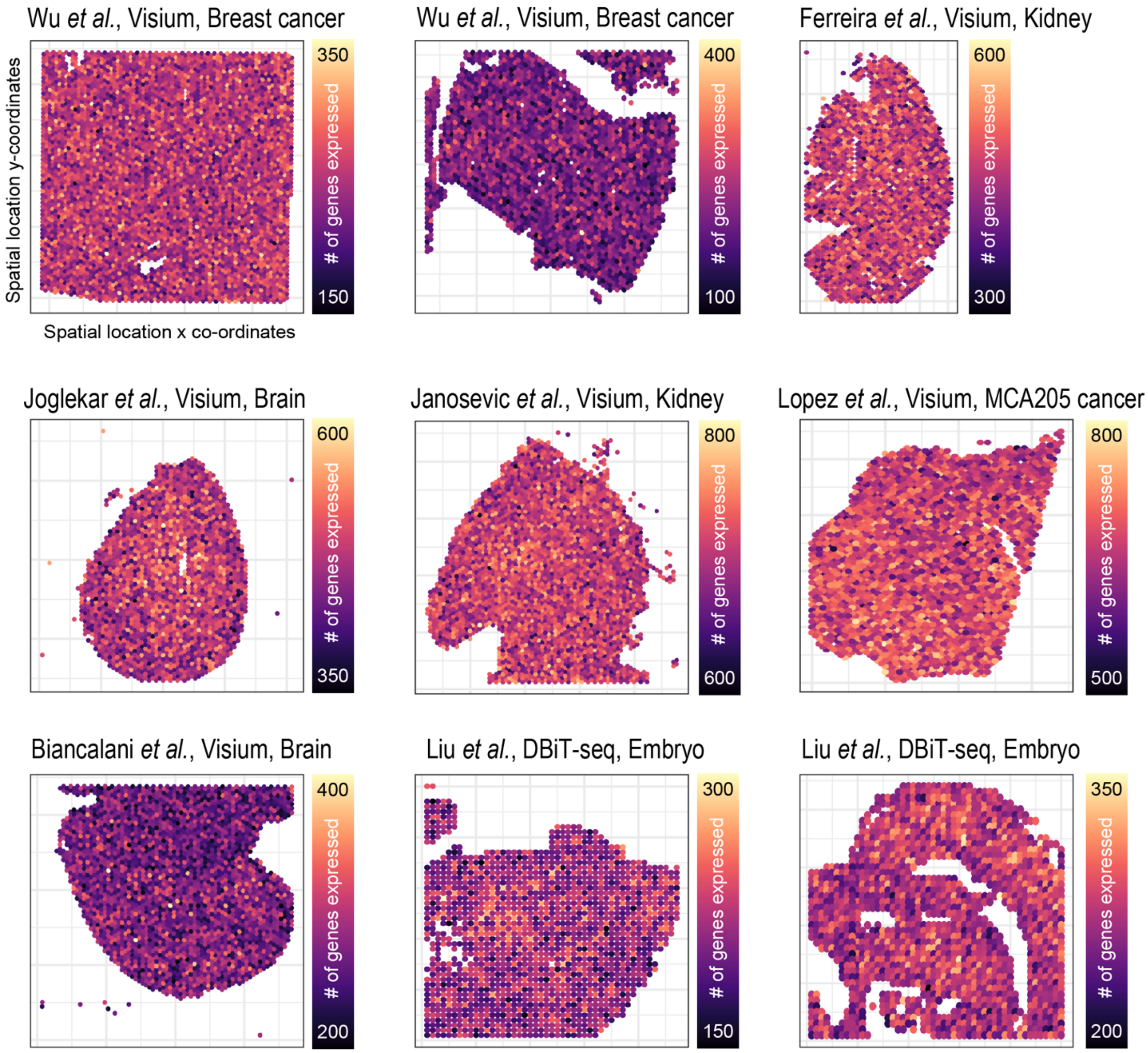


**Supplementary Figure 6. Simulation of spatial transcriptomics data.** Spatial locations and the total number of genes expressed per spatial spot in 9 simulated spatial transcriptomics datasets generated from different sources of data.

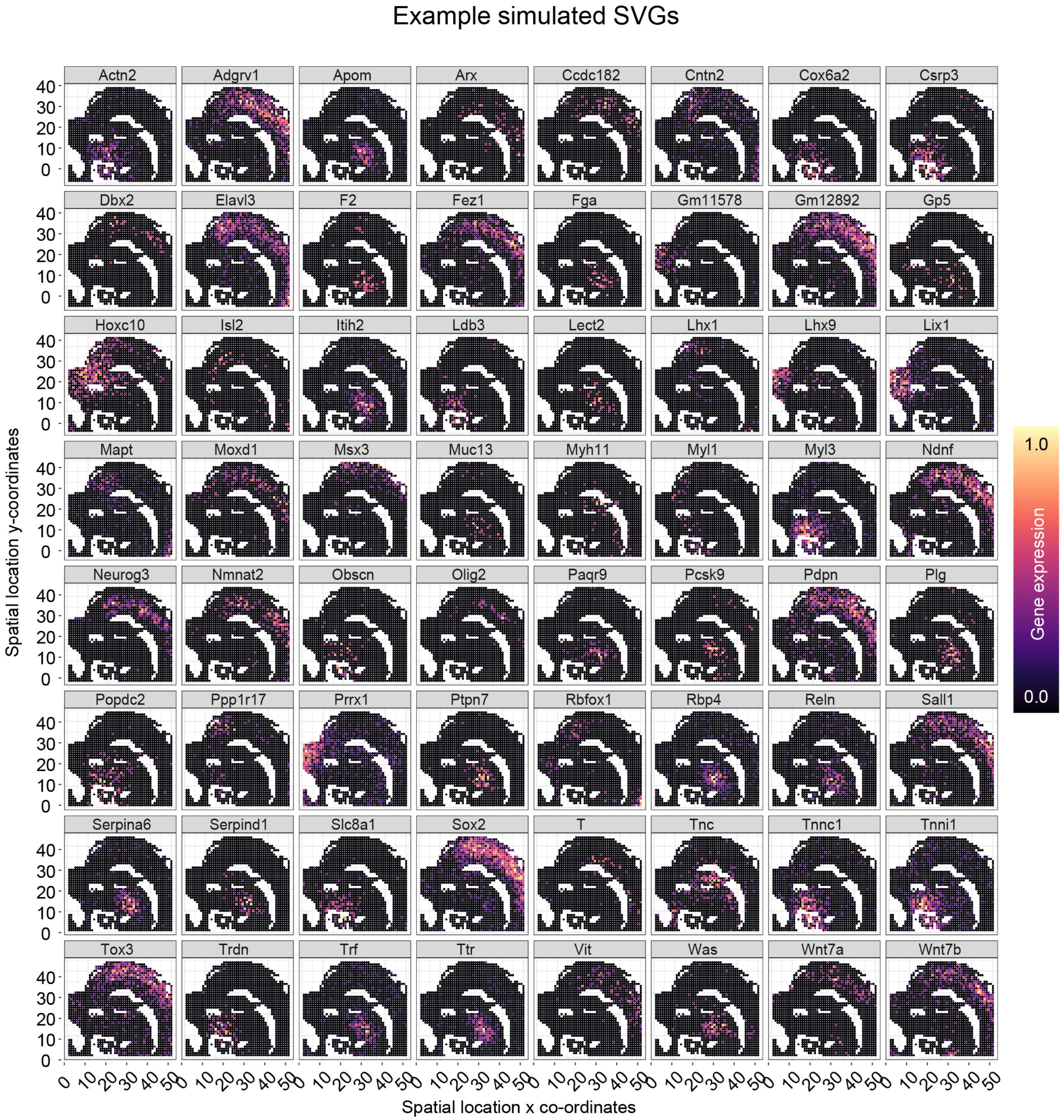


**Supplementary Figure 7. Spatial patterns of spatially variant genes.** Spatial expression patterns of example spatially variable genes of an example simulated spatial transcriptomics data (derived from Liu et al., DBiT-seq, Embryo).

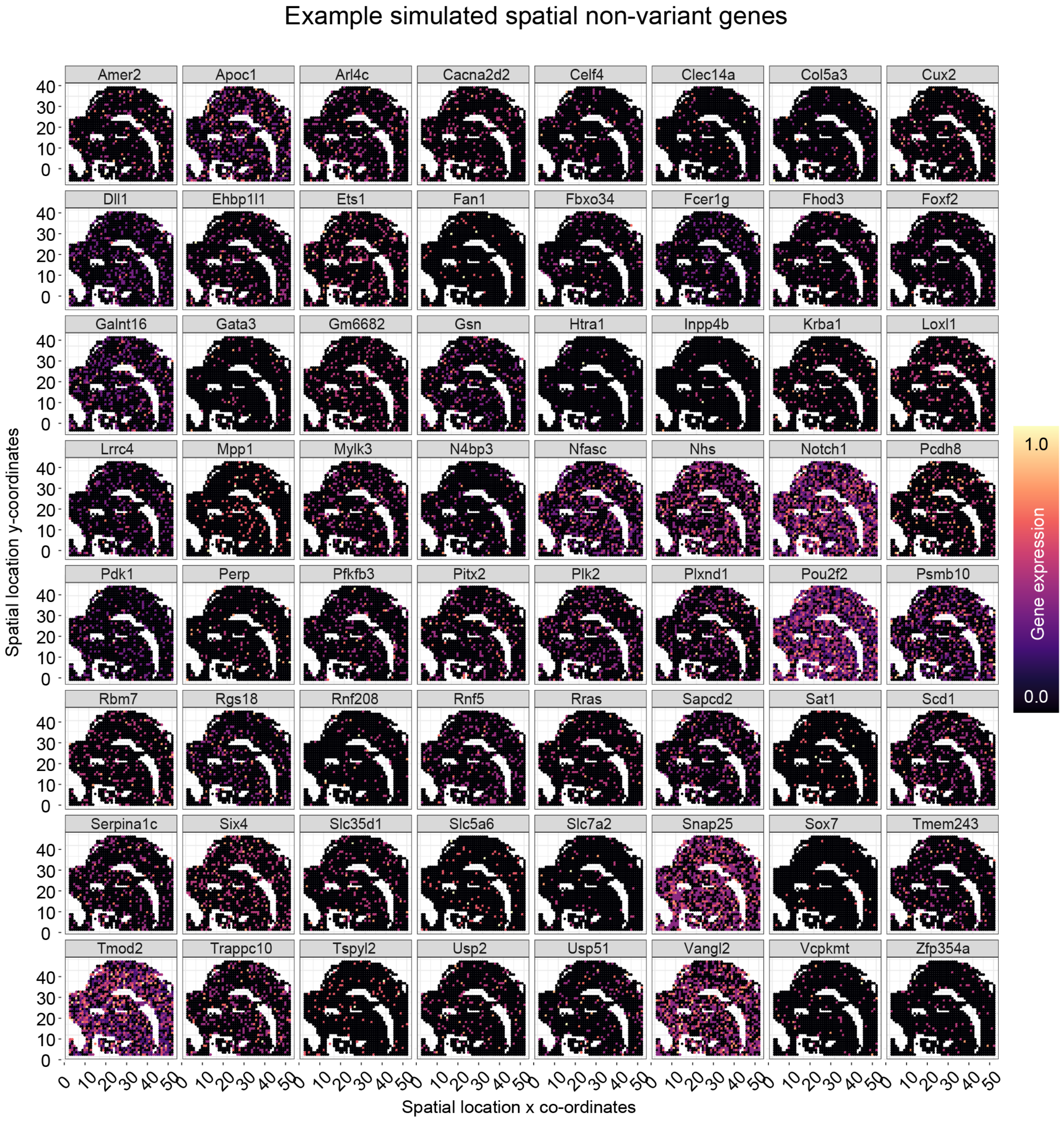


**Supplementary Figure 8. Spatial patterns of spatially invariant genes.** Spatial expression patterns of example non-spatially variable genes of an example simulated spatial transcriptomics data (derived from Liu et al., DBiT-seq, Embryo).

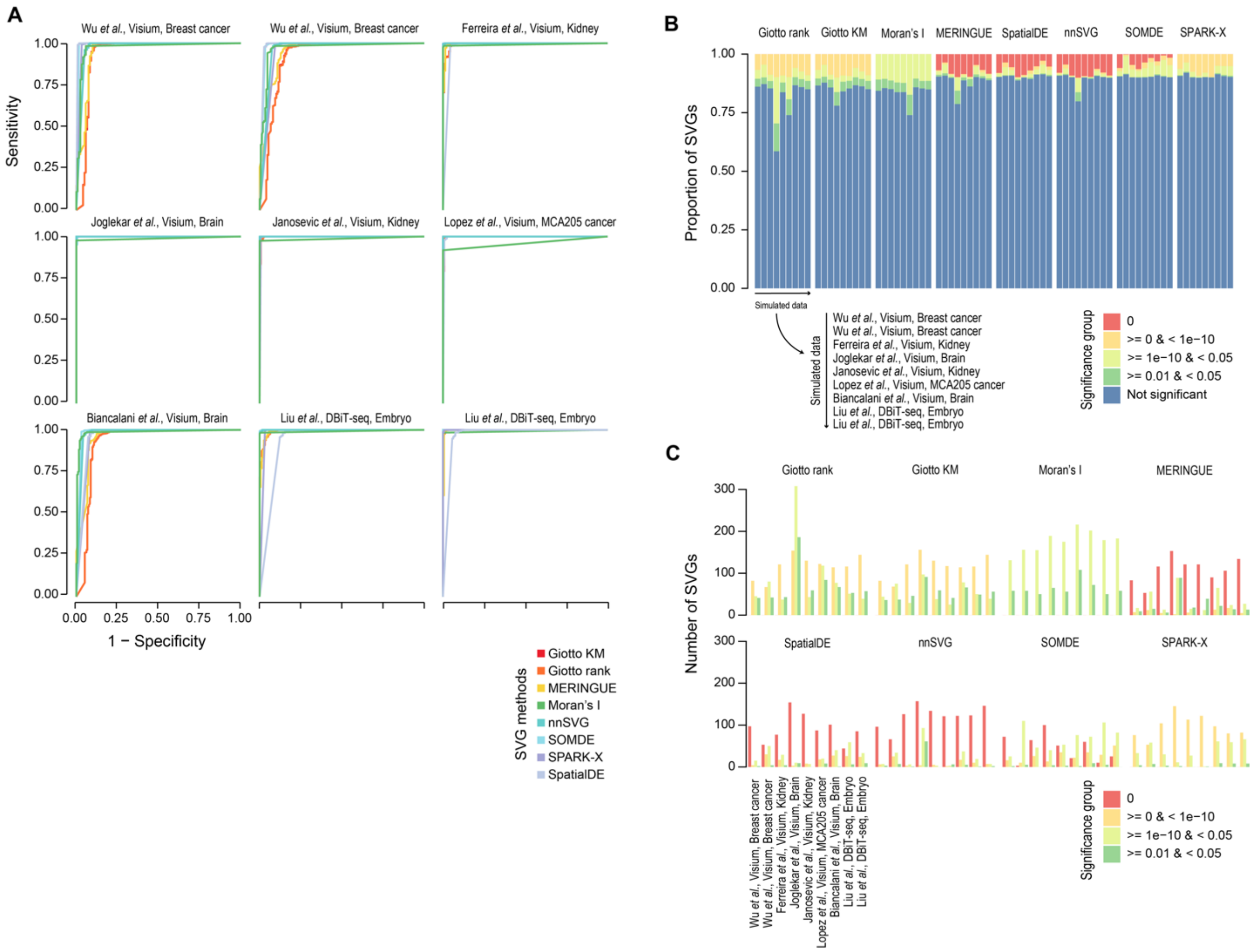


**Supplementary Figure 9. ROC curves of spatially variable gene detection. (a)** ROC curves were used to assess the capacity of each method to detect spatially variable genes between two spatial domains. The curves are colour-coded by methods. The 1-specificity and sensitivity of spatially variable gene detection are plotted on the x- and y-axes, respectively. **(b)** Proportion bar plots of SVGs of different statistical significance levels identified by each method. SVGs are partitioned into five categories based on the adjusted p-values reported by each method (i.e., p = 0; 0 < p <= 1e-10; 1e-10 < p <= 0.01; 0.01 < p <= 0.05; and p > 0.05) and presented as a percentage (y-axis). **(c)** Bar plots of the total number of SVGs reported by each method, for each simulated data, and for each statistical significance level as in **(b)**.

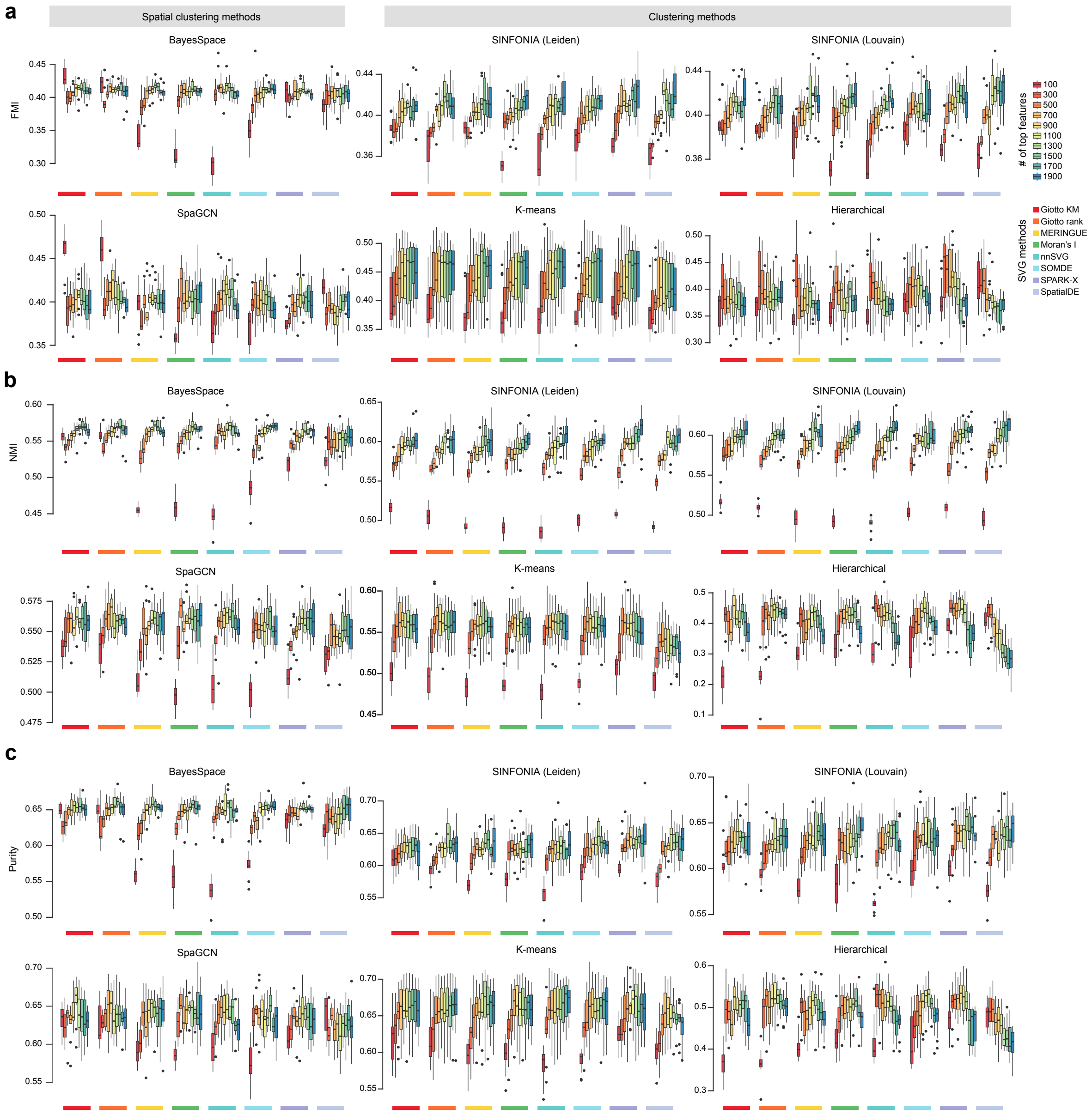


**Supplementary Figure 10. Performance of SVGs selected by each SVG method for clustering spatial domains in the mouse embryo.** Concordance in the clustering outputs from spatial clustering methods (BayesSpace and SpaGCN) and non-spatial clustering methods (SINFONIA's Louvain and Leiden, k-means, and hierarchical clustering) and the pre-defined spatial domains in the mouse embryo was computed using across a range of top SVGs (between 100 and 1900 genes). Concordance was quantified by three metrics including **(a)** Fowlkes-Mallows index (FMI), **(b)** normalized mutual information (NMI), and **(c)** purity.
